# Kronos scRT: a uniform framework for single-cell replication timing analysis

**DOI:** 10.1101/2021.09.01.458599

**Authors:** Stefano Gnan, Joseph M. Josephides, Xia Wu, Manuela Spagnuolo, Dalila Saulebekova, Mylène Bohec, Marie Dumont, Laura G. Baudrin, Daniele Fachinetti, Sylvain Baulande, Chun-Long Chen

## Abstract

Mammalian genomes are replicated in a cell-type specific order and in coordination with transcription and chromatin organization. Although the field of replication is also entering the single-cell era, current studies require cell sorting, individual cell processing and have yielded a limited number (<100) of cells. Here, we have developed Kronos scRT (https://github.com/CL-CHEN-Lab/Kronos_scRT), a software for single-cell Replication Timing (scRT) analysis. Kronos scRT does not require a specific platform nor cell sorting, allowing the investigation of large datasets obtained from asynchronous cells. Analysis of published available data and droplet-based scWGS data generated in the current study, allows exploitation of scRT data from thousands of cells for different mouse and human cell lines. Our results demonstrate that, although most cells replicate within a close timing range for a given genomic region, replication can also occur stochastically throughout S phase. Altogether, Kronos scRT allows investigating the RT program at a single-cell resolution for both homogeneous and heterogeneous cell populations in a fast and comprehensive manner.

## Introduction

DNA replication is a fundamental process in all living organisms that guarantees the duplication of genetic information before cell division. Based on inter-origin distance (∼one origin every 100 kb in mammalian cells) and replication fork speed (1-3 kb/min), if all replication origins were activated at the same time in a single mammalian cell, it would only take about 30 minutes for complete genome replication to occur^1,2^. However, due to limiting factors of replication initiation and replication fork progression, the DNA replication process is not simultaneously initiated at all the potential origins at once^3,4^. Rather, for each cell type, there is a defined selection and temporal order in which these origins fire, and thus, mammalian DNA replication takes several hours (usually between 6 and 12 h) to reach the finish line^5,6^. Furthermore, it has been shown that this temporal and spatial genomic orchestration coordinates with other processes such as chromatin organization and gene transcription^7–9^. The cell-type specific program that regulates the progression of DNA replication during the synthesis phase (S phase) is referred to as replication timing (RT) program^10,11^. It has been reported that RT is altered throughout disease development, such as in cancers and neurological disorders^12–14^. In addition, RT has been shown, by others and us, to play an important role in shaping the mutational landscape and impacting genome stability in both normal and cancer cells^15–19^. All these make RT an important aspect of better understanding the underlying causes, or outcoming effects, of genomic instability.

During the last decade, the advent of high-throughput single-cell omics has allowed the study of intercellular variability and has shed light on both functional and structural dynamics of cells. Advances in high-throughput single-cell sequencing techniques have also made it possible to analyze RT at the single-cell level. Recent single-cell RT (scRT) studies^20–23^, offer an advantage over bulk cell studies as they provide the possibility of studying the RT program within individual cells and enable the investigation of cell-to-cell variability. However, all current scRT studies require identification of cell phases (i.e. G1, S) at a pre-sequencing stage with fluorescence-activated cell sorting (FACS) and manual processing on plates of individual cells, allowing incorporation of only tens to a hundred cells^21–23^ which leads to scalability concerns. Furthermore, these analyses lack a unified pipeline to extrapolate scRT. Nevertheless, the advent of scRT investigations has supported the hypothesis that most replication domains follow the predetermined replication times as the majority of the other cells in the same population^21–23^. However, there is still room to explore non-conforming events in single cells that deviate from the average replication time and possibly follow stochastic replication. Importantly, latest advancements in Optical Replication Mapping (ORM) allow newly replicated DNA to be mapped and thus, early initiation events to be tracked at the single-molecule level^24^. Analysis of individual initiation events with ORM from human cells synchronized at the very beginning of S phase has revealed that, although most early initiation events occur in early-replicating regions of the genome, a significant number, ∼9%, occur in late replicating regions, supporting a stochastic model of initiation of replication. Therefore, the power of current scRT studies is shadowed by the previously mentioned limitations and the rare stochastic RT events may not be reflected in currently available scRT with limited sample sizes.

Here, to overcome current restrictions in scRT studies, we put forth a uniform computational framework named Kronos scRT to investigate scRT based on single-cell copy number variation (scCNV) detection from single-cell Whole Genome Sequencing (scWGS) data. Our pipeline can be used to analyze datasets from various experiments including classical single-cell Whole Genome Amplification (scWGA), droplet-based 10x Genomics Chromium scCNV solution, single-cell High-throughput Chromosome conformation capture (scHi-C) and other related data, obtained either from FACS sorted cells or directly from asynchronous cycling populations. The framework described here allows us to increase the number of cells used to analyze scRT by at least 10-fold (>1,000 cells in one experiment) compared to previous studies. By analyzing published data and the new droplet-based scWGS data generated in the current study, we obtained large amounts (up to 1,353 S-phase cells for a given cell type from a single experiment, 4,724 cells analyzed in total) of scRT data of different mammalian cell lines. The obtained scRT data allow us to construct distinguished S-phase progression trajectories of different cell types as well as to identify coexisting sub-populations. In addition, our results show that incorporating significantly more cells enables us to stretch the study of DNA replication heterogeneity with unprecedent detail. Our analysis demonstrates that, for given genomic regions, although most cells replicate within a close timing range around the population average RT, replication can also happen stochastically in any given window during S phase. Modeling of replication kinetics by using scRT data demonstrates that measuring the firing efficiency in early S phase can predict the average firing time within a cell population. Here, we extent a previous analysis from the single-molecule level into the single-cell level and show that stochastic regulation of replication kinetics is a fundamental feature of eukaryotic replication.

## Results

### Kronos scRT: a computational tool allowing extraction and analysis of scRT data from scWGS data of asynchronous cells

We have developed Kronos scRT, a tool that computes scRT under a unified framework and in a comprehensive manner (Fig. 1a and Supplementary Fig. 1a). At first, single-cell DNA sequencing reads are aligned to the provided reference genome and counted over regular bins (by default, and in our analysis 20 kb), of which the size can be adjusted depending on the average coverage of the experiment. Read counts are then corrected for GC content and mappability bias. An option to blacklist genomic regions is available as well (Methods). The data are then segmented and copy number variation (CNV) is called independently within each individual cell. At this stage, two additional parameters are calculated: the cell ploidy, as the cell weighted mean copy number (CN), and the intracellular bin-to-bin variability, calculated as Depth Independent Median Absolute deviation of Pairwise Differences (DIMAPD) (Methods). The following step consists of the identification of G1/G2-phase and S-phase cells. Depending on the type of data and the information available, Kronos scRT encompasses different approaches to distinguish cells in G1/G2 phase from those in S phase. If cells are FACS sorted in discrete populations, as in previous papers^21–23^, the phase information can be used directly to label cells into these two groups. Otherwise, for unsorted cycling populations, the detection of the S-phase cells can be calculated automatically. The automatic detection is based on two assumptions: firstly, most cells belong to the G1/G2 population; secondly, the intracellular bin-to-bin variability is minimal in G1 and G2 cells, where all the bins have similar CN, and maximizes towards mid-S phase due to the asynchronous replication of adjacent bins (Fig. 1b,c). The program, therefore, fits the variability data into a gaussian distribution and identifies S-phase cells as outliers (Methods). Lastly, if the cell cycle distribution has been altered, i.e. cells have been enriched for the S-phase without complete removal of G1 and G2 phase cells, the user can manually impose a variability threshold based on visual inspection of the data as in Fig. 1b.

**Figure 1.**
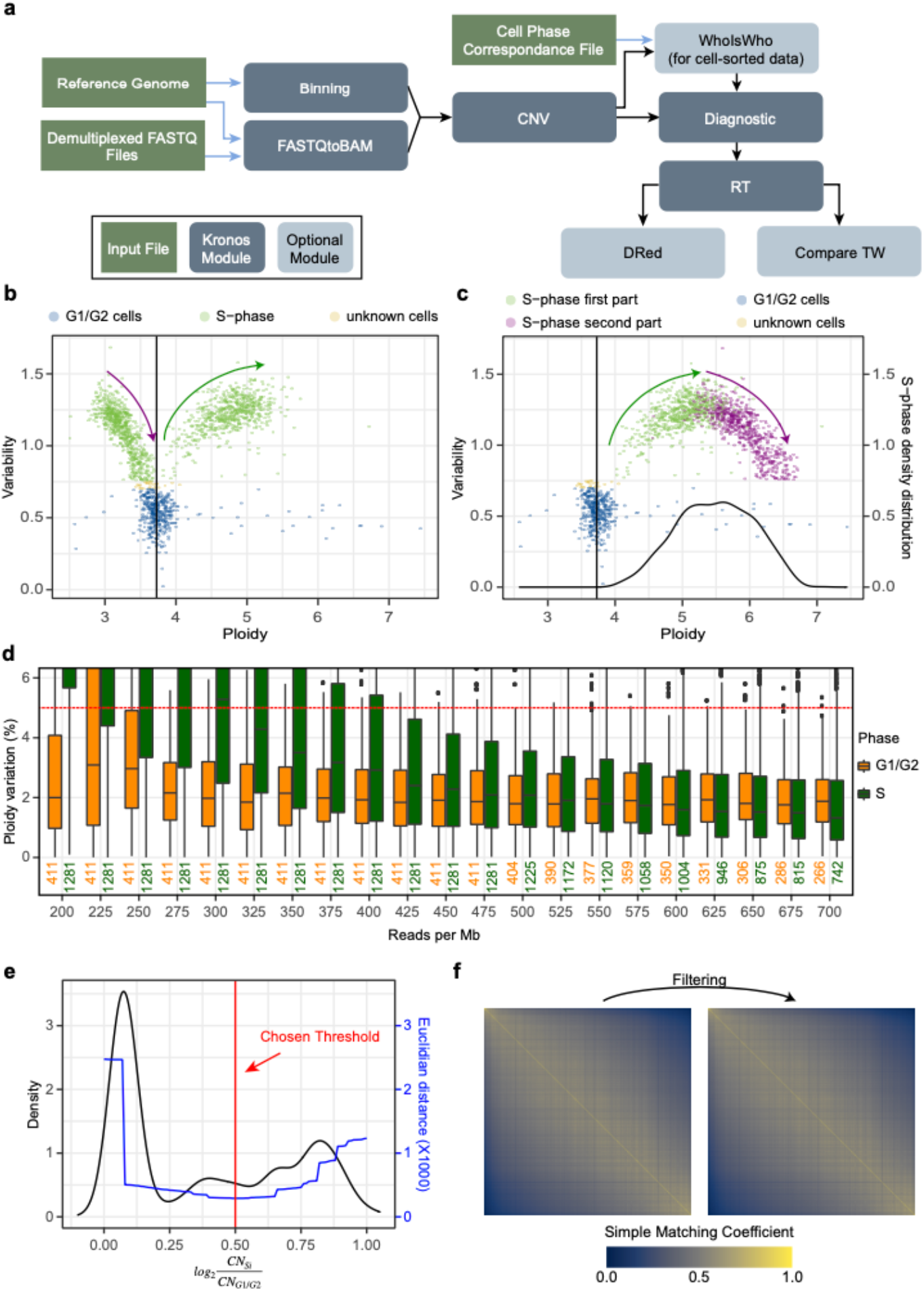
An efficient uniform framework for scRT extraction. **a** The pipeline of Kronos scRT with the different modules enlisted. The input files, the main modules of Kronos RT and the optional modules are shown in green, dark blue and light blue, respectively. **b** Scatter plot reporting the average ploidy of a cell on the x-axis and the bin-to-bin intracellular variability on the y-axis. Each point is a single cell and the color is assigned based on a cut-off automatically calculated, or manually imposed, regarding the population variability (Methods). High variability is associated with S-phase cells (green), while low variability is associated with G1/G2-phase cells (blue). An unknown region can be manually set (yellow). The vertical black line represents the median ploidy of the population. Reported data are from asynchronized MCF7 cells enriched for the S phase. The green and purple arrows show the S-phase progression of the first and second part of the S phase, respectively (Methods). **c** Data presented in (b) after S-phase progression correction. The color of the second portion of the S phase have been changed to purple. As in (b), the green and the purple arrows indicate the correct S-phase progression. The black curve reports the S-phase density distribution that is used to calculate the parameters to adjust the S-phase progression (Methods). **d** Read down-sampling for the dataset of MCF7 cells obtained by the 10x Genomics system. G1/G2- and S-phase cells are plotted in orange and green, respectively. Boxplots report the percentage change of the mean ploidy after down-sampling compared to the original value. The red dashed line indicates a change of 5%. Cells with more than 450 reads per Mb were used for the down-sampling, and the number of cells used for each down-sampling is reported below the boxplot. **e** A normalized copy-number (nCN) distribution of a representative single cell in black. In blue, the Euclidian distance between the real data and a binarized system computed with a certain nCN cut-off. In red, the cut-off used to binarized the data that corresponds to lowest calculated Euclidian distance (Methods). The nCN of a bin is calculated as the log2 ratio of the CN of that bin in a replicating cell and the median CN of the same bin in the non-replicating population. **f** Simple matching coefficient matrix among single cells. The identified irregular cells are therefore discarded as a further measure of quality control (see Methods for details).

Due to the way of calculating the CN (Methods), it is impossible to discriminate between G1 and G2 cells. This affects the S-phase population as well, which is split into two parts. The first part has higher ploidy than the G1/G2-phase population and shows increasing variability moving towards higher ploidy. The second part, instead, shows a lower ploidy than the G1/G2 pool and its variability decreases while approaching G1/G2 population (Fig. 1b). S-phase cell ploidy must therefore be adjusted before proceeding with the downstream analysis. Adjustment can be automatically calculated (Fig. 1c) or manually imposed by the user (Methods).

The cells shown in Figure 1b-d come from a MCF7 breast cancer cell line dataset created using 10x Genomics microfluidic system. According to ATCC (American Type Culture Collection), this cell line has 80 autosomes in G1 (ploidy: 3.64), which agrees with the estimation of Kronos scRT (Fig. 1b-c and Supplementary Fig. 3b). In addition to the ploidy estimation step, cells with a low coverage are filtered out. To evaluate the minimum number of reads required to obtain a stable CNV calling with each experimental setting, we selected G1/G2- and S-phase cells with relative high coverages and down-sampled them. Based on this simulation, the ploidy estimations of G1/G2-phase cells are quite stable while S-phase cell mean ploidy calling is more sensitive to coverage changes (Fig. 1d and Supplementary Fig. 1b,c). As a threshold, we selected the smallest down-sampling value having 75% of S-phase cells with a ploidy estimation within 5% variation from the original one (Fig. 1d and Supplementary Fig. 1b,c). Therefore, for our MCF7 cells, we can estimate that a minimum of 117 reads per megabase (RPMb) per haploid genome (425 RPMb / 3.64 ploidy) is needed (i.e. about 0.75 million reads for a diploid human genome) in order to have a stable mean ploidy estimation.

Adjusted CNs of cells that passed the filter can then be used to calculate scRT profiles. Based on the coverage of our data, we binned the genome into 200 kb non-overlapping windows and calculated the weighted median CN for each cell. Using the G1/G2-phase population, a median CN profile is calculated and used to normalize the CN of each S-phase cell as a log2 ratio (Methods). Data are then binarized to obtain the scRT profiles, where 1 corresponds to replicated regions and 0 to unreplicated ones. This is based on the assumption that replicated regions will have a doubled CN compared to the G1/G2 fraction, and therefore a log2 ratio close to 1, while non-replicated regions will have the same CN and a log2 ratio close to 0. Binarization is independent for each cell and is based on the identification of a normalized CN threshold. To identify the most appropriate threshold, we select the one that minimizes the Euclidian distance between the generated scRT profiles and the original data (Methods). An example is shown in Fig. 1e. As an additional quality control, a pairwise Simple Matching Coefficient of the scRT profiles is calculated and cells with a RT profile deviating from the main population are filtered out (Fig. 1f) (Methods). Finally, we can compute the pseudo-bulk RT as the weighted mean of the scRT profiles and compare it with the bulk RT.

In conclusion, by combining CNV calling, inter-cellular variability and quality control filtering, we successfully established a unified and genome-wide scRT computational profiling tool, Kronos scRT, which allows studying scRT from scWGS of both FACS sorted and unsorted cycling cells in an efficient manner.

### Determination and analysis of scRT from scWGS data of sorted S-phase cells

scRT studies usually require cell phase sorting via FACS because they lack of an *in silico* phase separation as presented in the current study. Nevertheless, cell phase information can be integrated to the analysis to label cells using the WhoIsWho module (Fig. 1a). To test the applicability of Kronos scRT’s framework, we used previously published scWGS data derived from sorted mid-S-phase mouse embryonic stem cells (mESC, n=67 cells) and mESC differentiated for 2 days to EpiLCs followed by five days of Embryo Bodies culture, which result into neuroectoderm cells (hereafter called NE-7d, n=45 cells)^22^. The scRT profiles of these two cell types were determined using Kronos scRT, and the phase was assigned using the FACS metadata (Supplementary Table 1 and 2).

To demonstrate that Kronos scRT is efficient in detecting scRT profiles, even with the utilization of a small number of sorted mid-S-phase cells, we calculated the correlation between pseudo-bulk RT issued from scRT of each sample in the current study and BrdU-IP-issued bulk RT data generated by Takahashi and colleagues^22^ (Fig. 2a). We obtained a Spearman correlation of 0.890 for mESC and 0.902 for NE-7d cells (Fig. 2b and Supplementary Fig. 2a), demonstrating the robustness of our method and computational pipeline. In addition, in agreement with previous studies^21–23^, the obtained mid-S binary replication signals show that cell-to-cell variability exists but is limited and that RT organization is largely conserved in single-cells.

**Figure 2.**
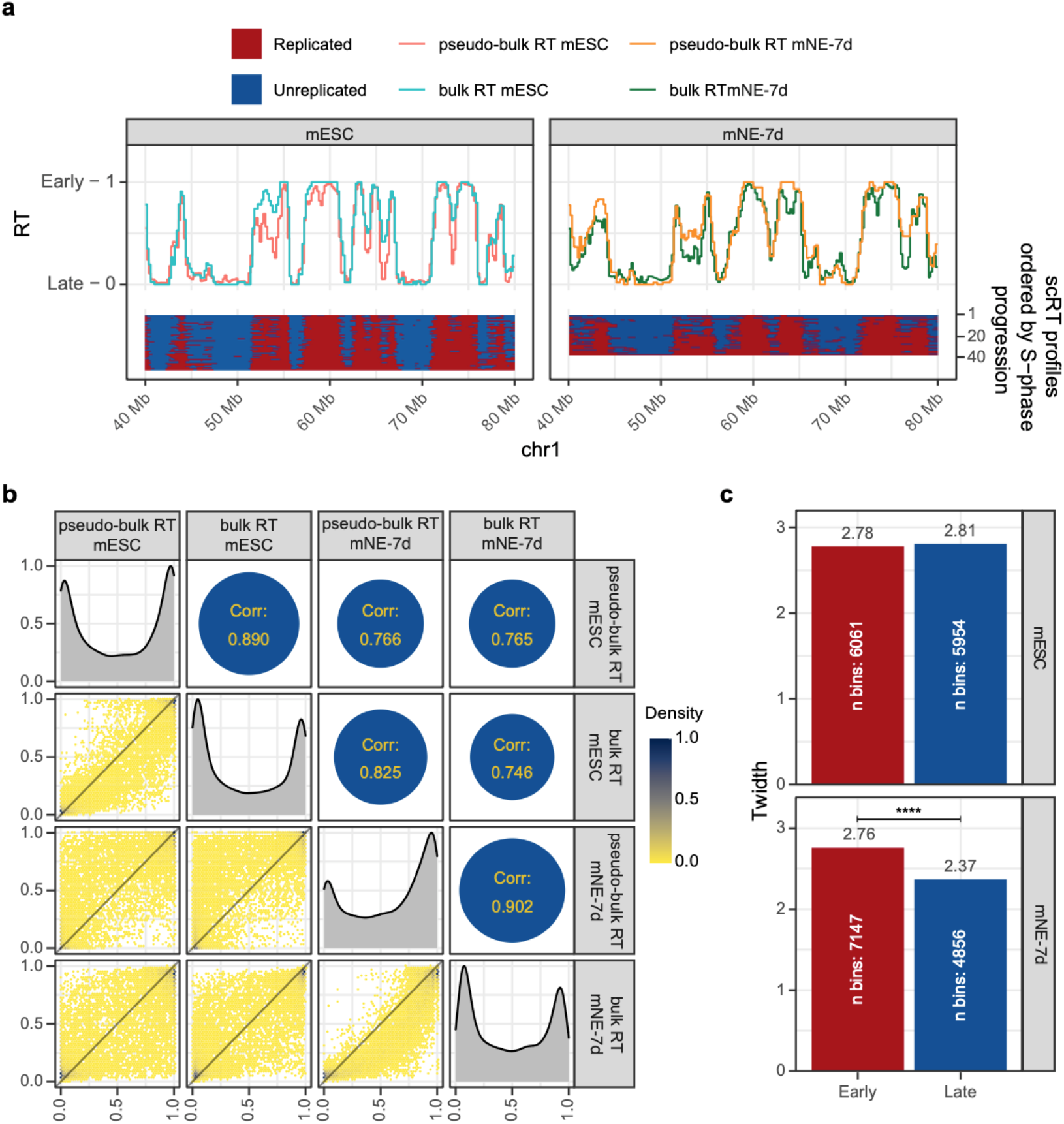
Extracting scRT from sorted mid-S phase mouse cells. **a** scRT calculated from FACS sorted mid-S phase mouse embryonic stem cells (mESC, left) and 7-day differentiated neuroectoderm cells (mNE-7d, right). In the bottom panel: the binary scRT profiles are reported sorting single cells from top to bottom by increasing replication percentage. The replicated and unreplicated regions are shown in red and blue, respectively. In the upper panel, the pseudo-bulk RT calculated from the scRT profiles (see Methods) are compared to the bulk RT of the corresponding cell type. **b** Comparison between single-cell and bulk RT data. Lower-triangle: 2D density plot reporting pair-wise comparisons between samples. Density color code is reported on the right. Upper-triangle: Spearman correlation between RT data. Diagonal: RT distribution of each sample. **c** T_width_ values of mESC (upper) and mNE-7d (lower) for Early (RT > 0.5) and Late (RT ≤ 0.5) replicating regions. Higher T_width_ indicates higher RT variability among cells. P-values were calculated using the Kronos scRT Compare TW module (see Methods, **** < 10^−4^)

We were then interested in quantifying the replication variability within the cell populations by calculating, in each dataset, the T_width_ values, defined as the time needed for a given genomic region to be replicated from 25% to 75% of cells in a S-phase lasting 10 h^21^. We found that the T_width_ in mESCs ranged between 2.78 h for early and 2.81 h for late replicating regions, while, in mNE-7d cells, it ranged between 2.76 h for early and 2.37 h for late. Using the Compare TW module, we applied a null hypothesis test through bootstrapping with H_0_: T_width_early_ = T_width_late_ and with H_1_: T_width_early_ ≠ T_width_late_ (Fig. 2c). No significant statistical difference can be observed between early and late T_width_ (p-val=0.43) for the mESC cells, while for the mNE-7d cells, late replicating regions seem to be less variable than early ones (p-val < 10^−4^). Based on these results, mESC present a behavior similar to what was observed in Dileep & Gilbert 2018^21^, with no significant differences between early and late S-phase, while mNE-7d shows lower variability at the end of the S-phase.

### Determination of scRT data from asynchronous cells using a microfluidic-based system

To demonstrate that Kronos scRT can detect scRT without cell sorting and cell phase identification (e.g. by FACS), we first generated scWGS data using a droplet-based 10x Genomics Chromium scCNV solution for 368 estrogen-treated cycling MCF7 cells (containing about 20% of S-phase cells) (Fig. 3a) (see Methods) (Supplementary Table 1). As previously discussed, this condition allowed us to use the automatic identification of the S-phase cells (Fig. 3a, left panel). Moreover, the even distribution of cells across the S phase allowed the usage of the automatic S-phase correction (Fig. 3a, right panel). We calculated the scRT and pseudo-bulk from 82 identified S-phase cells (Fig. 3b and Supplementary Fig. 2b). The pseudo-bulk RT shows a high degree of correlation (Spearman correlation R=0.913) with the bulk RT (Fig. 3c). By visual inspection of the scRT profiles, it was possible to clearly identify a certain degree of variability (Fig. 3b). An advantage of having cells evenly distributed throughout S phase is the possibility to perform the T_width_ analysis with higher resolution than only with the mid-S-phase-sorted cells (Fig. 3d). As suggested in previous studies^22,25^, T_width_ values are smaller at the beginning and at the end of the S phase, and with a progressive increase while moving towards the mid-S phase. Although with a limited number of observations (only 82 cells), such results reinforce the idea that initiation and termination of the replication program are more tightly regulated compared to the mid-S phase (Fig. 3d).

**Figure 3.**
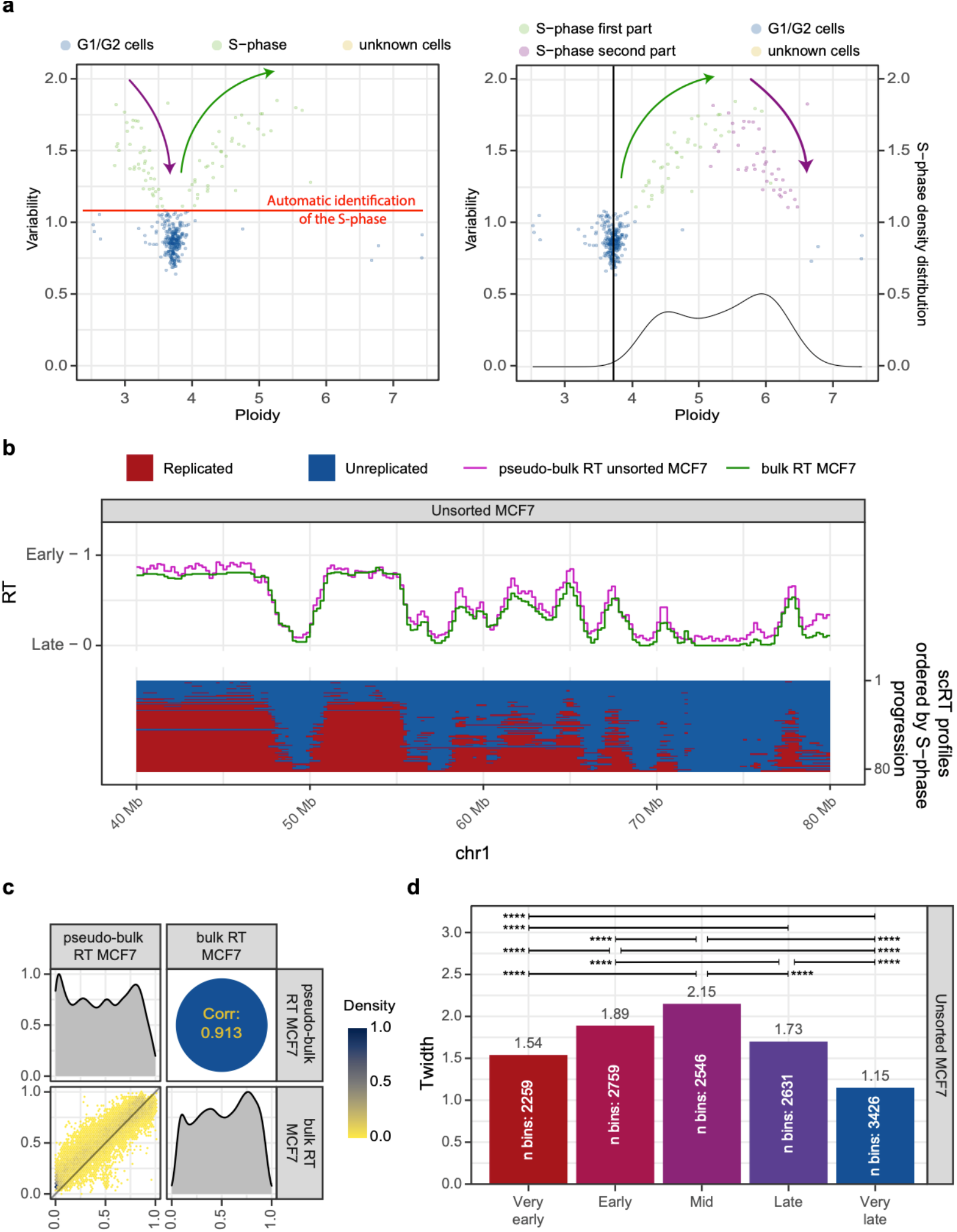
Extracting scRT from asynchronous human MCF7 cells. **a** Intracellular bin-to-bin variability scatter plot in function of single-cell ploidy before (on the left) and after (on the right) S-phase-progression correction. The color code is the same as Fig. 1b and c. Kronos scRT is able to automatically identify the S-phase cells from the asynchronous scWGS data (see Methods). **b** scRT calculated from unsorted cycling MCF7 breast cancer cells. Same presentation as in Fig. 2a **c** Comparison between single-cell and bulk RT data, as in Fig. 2b. **d** Barplots reporting the T_width_ calculated on regions of 5 RT categories based on the pseudo-bulk RT values (Very early > 0.8, 0.8 ≥ Early > 0.6, 0.6 ≥ Mid > 0.4, 0.4 ≥ Late > 0.2, Very Late ≤ 0.2). P-values were calculated using the Kronos scRT Compare TW module (Methods, * < 0.05, ** < 10^−2^, *** < 10^−3^, **** < 10^−4^).

### scCNV/scRT analyses allow identifying sub-populations of cells within a heterogeneous population

To obtain a more representative estimation of cell-to-cell replication timing variability, we decided to further increase the number of S-phase cells. As unsorted cell populations generally contain many more G1/G2-than S-phase cells, to reduce the sequencing costs, we enriched these samples for the S-phase through FACS sorting using a gate that contains majority the S-phase cells (Methods) following with scWGS. Using this approach, we obtained 1,777 MCF7 S-phase enriched cells, the majority of which (n=1,353) belong to S-phase cells (Fig. 1b,c, Supplementary Table 1).

While preforming a dimensionality reduction analysis with Uniform Manifold Approximation and Projection (UMAP)^26,27^, on the scRT profiles of MCF7 S-phase enriched cells, we noticed that cells were disposed in an arch arrangement (Supplementary Fig. 3a). After taking into consideration percentage of replication of each cell, we realized that the arch was due to the presence of at least two different replication timing groups whose similarity is maximised at the beginning of the S phase (most of the genome in both groups is 0) (Supplementary Fig. 3a). We reasoned that the presence of two replication timing groups, in this cell line, would most probably be associated with two sub-populations of cells with distinguish chromosomal rearrangements. We, therefore, reperformed the dimensionality reduction analysis using the scCNV instead of the scRT from both G1/G2- and S-phase cells (Fig. 4a). This analysis showed a clear separation of 4 groups: two G1/G2-phase groups and two S-phase ones (Fig. 4a). While the average G1/G2 ploidy of the MCF7 population was around 3.73, sub-population 1 had a ploidy of 3.68 and sup-population 2 had ploidy 3.84 (Supplementary Fig. 3b). As expected, CNV differences were observed between the two sub-populations, with the majority involving chromosomes 3, 7, 8, 11, 18 and 19 (Fig. 4b and Supplementary Fig. 3c). Chromosome 3 was of major interest as the copy-number profiles suggested a clear chromosome-wide copy-number gain (Fig. 4b and Supplementary Fig. 3c) in sub-population 2 compared to sub-population 1. To further confirm this copy-number gain, we performed Fluorescence *in situ* Hybridization (FISH) on metaphase spreads with a probe specific for the entire chromosome 3 and one for its centromere (Fig. 4c,d). We found that most cells had 4 or 5 copies of chromosome 3, with respectively 162 (47.09%) and 170 (49.42%) out of 344 analyzed spreads (Fig. 4c-e), in agreement with our observation on scCNV data (Fig. 4b and Supplementary Fig. 3c). We were, therefore, able to associate each G1/G2-phase group to its corresponding S-phase group, and pursue our analysis using the correct normalization for the data and deduce scRT of each sub-population of MCF7 cells individually. Our analysis demonstrates that Kronos scRT can analyze replication timing program within heterogeneous cell cultures and identify the underling sub-populations, which cannot be achieved with current population-based approaches.

**Figure 4.**
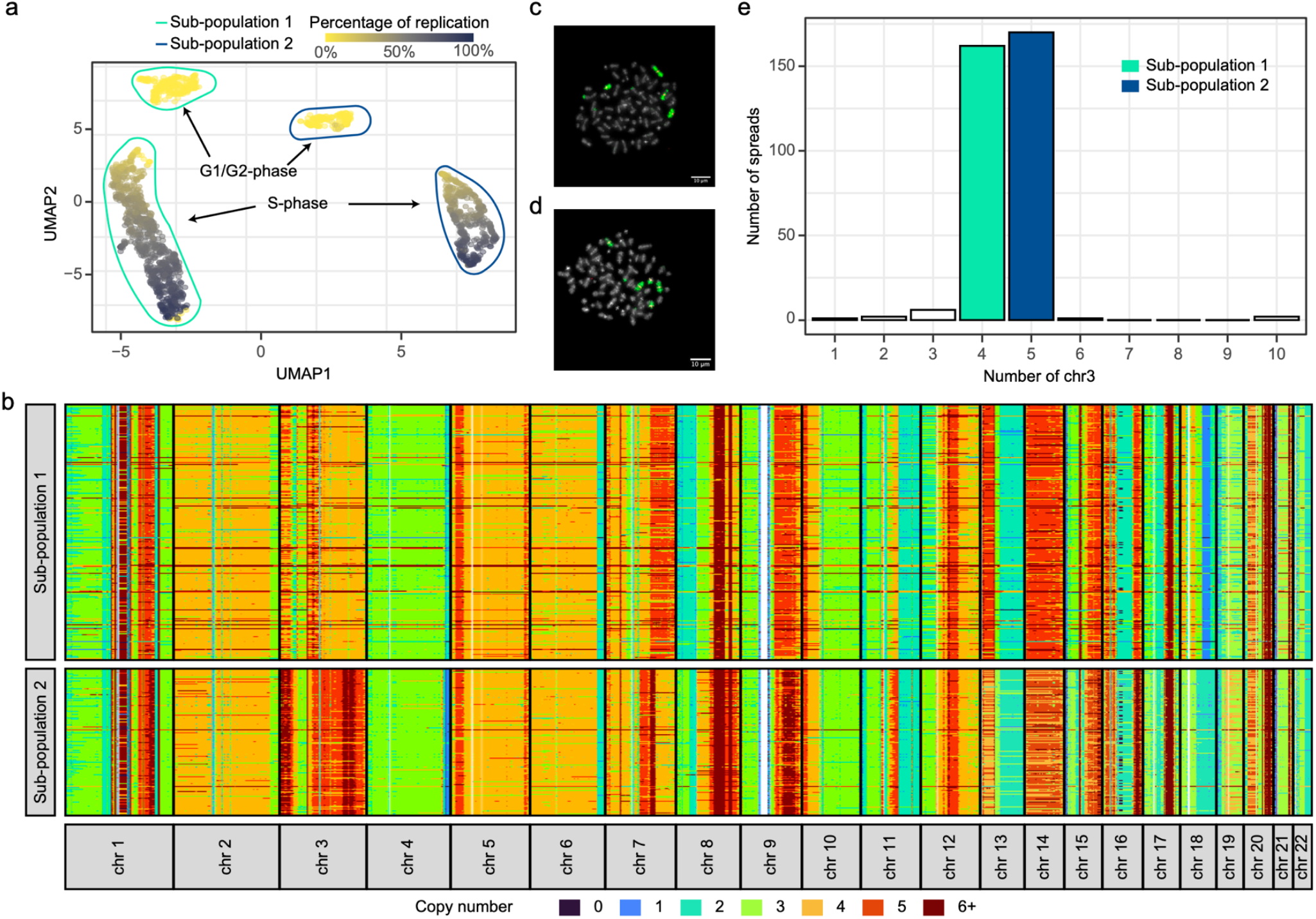
Identification of two sub-populations in MCF7 cell culture. **a** UMAP on scCNV of MCF7 cells colored by S-phase progression (only autosomal chromosomes). Both G1/G2 (yellow) and S-phase (gradient colored as indicated) cells are separated in 2 major sub-populations. G1/G2- and S-phase cells are gated with color coded gates, with the sub-population 1 in aqua and the sub-population 2 in blue. **b** Copy-numbers along autosomes detected in G1/G2 MCF7 cells, which are separated into 2 sub-populations based on the clustering results shown in (a). The binning for the visualization is 1 Mb (median CN profile in Supplementary Fig. 3c). **c-d** Examples of FISH experiment for cells with 4 (**c**) and 5 (**d**) copies of chromosome 3 that correspond to the sub-population 1 and the sub-population 2, respectively. Chromosomes are stained with DAPI (grey), chromosome 3 is labelled in green and its centromere in red. Scale bar: 10 μm. **e** Chromosome 3 counts based on FISH images (n=344). The two major groups with 4 (n=162) and 5 (n=170) copies of chromosome 3 correspond to the two groups shown in (a), respectively sub-population 1 and sub-population 2.

### Determination of scRT spanning thousands of single cells of various human cell types

In addition to MCF7 cells, we further extended our analysis to other cancer cells, 514 HeLa cells (including 259 S-phase cells), and 1,106 Jeff cells (normal lymphoblastoid, including 960 S-phase cells) (Supplementary Table 1). Both MCF7 sub-populations were analyzed as an individual sample. Although their pseudo-bulk RT were extremely close to each other (R=0.946) (Fig. 5a,b), we were still able to distinguish two different sub-populations when preforming dimension reduction (Fig. 5c,d). Concerning the other two cell lines, we identified only one population with a median ploidy of 2.87 for HeLa cells, as well as one population with a median ploidy of 1.94 for the Jeff cells (Supplementary Fig. 3b). As for MCF7 cells, we calculated the scRT and pseudo-bulk RT profiles for HeLa and Jeff cells (Supplementary Fig. 4a,d). The pseudo-bulk RT highly correlates with the bulk RT of the corresponding cell types (Supplementary Fig. 2b and Supplementary Fig. 4b,e). The scRT profiles are unique for each cell type as well and can be separated through dimension reduction (Fig. 5c,d). Finally, we calculated the T_width_ using the same 5 classes of RT categories as in Fig. 3d. For all analyzed cell types, the regions replicated at the very beginning or very end of the S-phase are better synchronized, i.e. with lower T_width_ values ranged between 1.2-1.4 h in early and 1.0-1.2 h in late replicating regions, compared to regions replicated around mid-S phase, which instead have T_width_ values around 1.7-1.8 h (Fig. 5e and Supplementary Fig. 4c,f).

**Figure 5.**
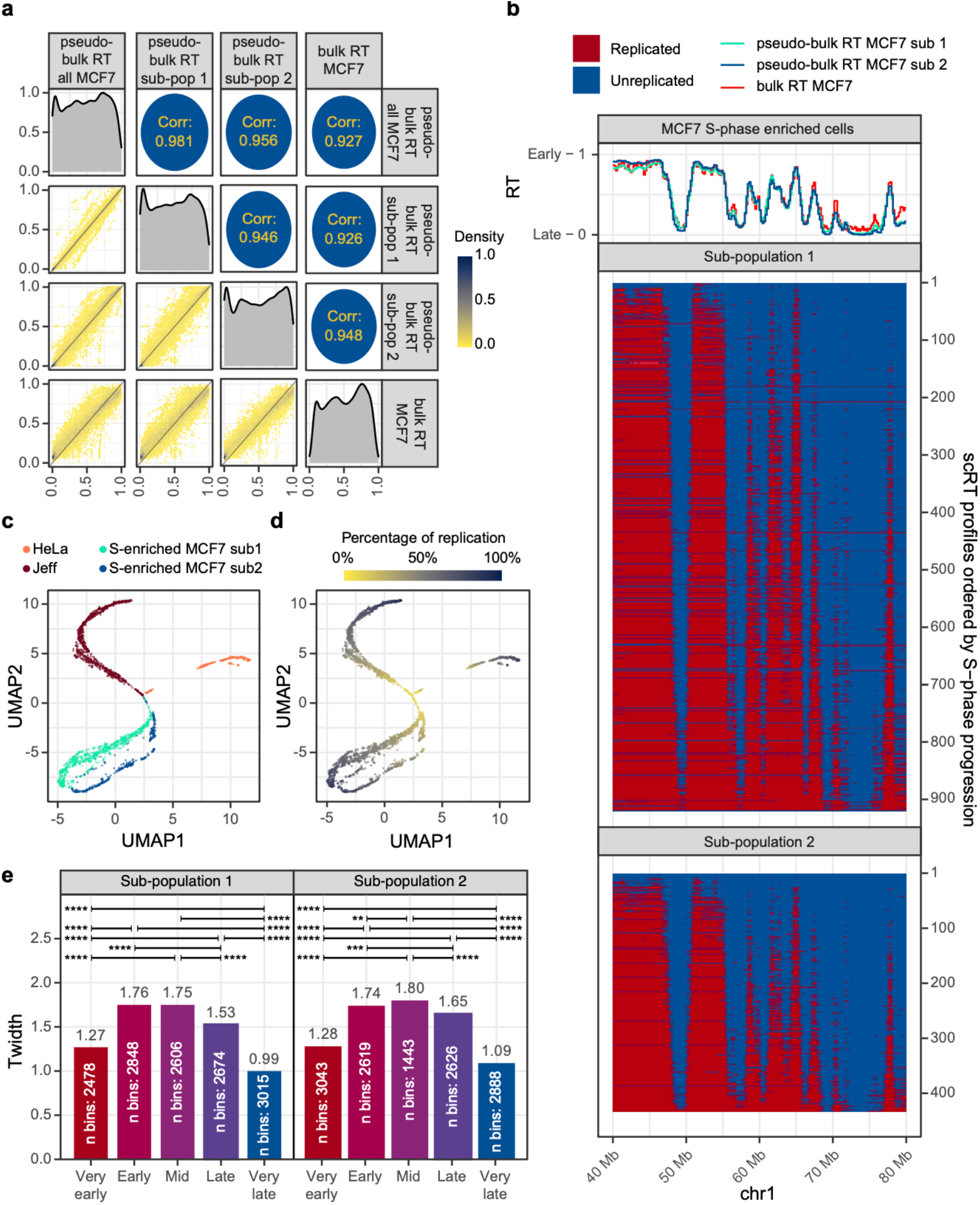
scRT from S-phase enriched human cells. **a** Pairwise comparison of the MCF7 pseudo-bulk RT and bulk RT profiles. Same as in Fig. 2b. **b** The scRT of S-phase enriched MCF7 cells over a representative region. In the upper part of the plot pseudo-bulk RT profiles of the two MCF7 sub-populations and bulk RT profile. In the bottom panel, the scRT profiles ordered from top to bottom by replication percentage of each cell. **c-d** Dimensionality reduction analysis of scRT data of different human cell types generated in the current study. Cells are color-coded based on cell type in (c) and based on replication percentage in (d). **e** Barplots reporting the T_widths_ calculated on 5 RT categories based on the pseudo-bulk RT values in the two MCF7 sub-populations. Categories were selected as in Fig. 3d. P-values were calculated using the Kronos scRT Compare TW module (see Methods, * < 0.05, ** < 10^−2^, *** < 10^−3^, **** < 10^−4^).

### scRT of human cells show a stochastic variation within a cell population

One of the main questions in the RT field is to elucidate whether this program is stochastic or deterministic. The authors of previous scRT studies commented on the improbability of the system to be stochastic since they reported that cell-to-cell variability is low^21–23^. However, their observations were based on limited number of cells and were therefore not able to identify rare events.

Recently, new evidence based on Optical Replication Mapping (ORM) sheds new light on this point^24,28^. Taking advantage of the high number of scRT obtained in the current study, we aimed to tackle this issue. To better visualize stochasticity, we selected the S-phase cells from 3 representative stages: early-S-phase cells (≤ 30% replication), mid-S-phase cells (40-60%) and late-S-phase cells (≥ 70%). We then assigned each 200 kb genomic bin to an RT category based on its pseudo-bulk RT, and finally, for each bin, we calculated its probability of being replicated in each representative stage (Fig. 6a,b and Supplementary Fig. 5a,b). If the RT program was deterministic, we would not expect to see late replicating regions (RT < 0.5) being replicated in the early-S-phase cells, and *vice versa*. As expected, for the mid-S-phase cells, the majority of early replicating regions are replicated, while most late replicating regions are unreplicated (Fig. 6a,b and Supplementary Fig. 5a,b, middle panel). Strikingly, in all the examined cell lines, we can observe 1-5% of cells replicating late genomic domains (pseudo-bulk RT < 0.5) at the beginning of the S-phase (cells with less than 30% replication) (Fig. 6a,b and Supplementary Fig. 5a,b, left panel). At the same time, if we focus on the late-S-phase cells (cells with more than 70% replication), in the deterministic model, we would expect all early replicating regions (pseudo-bulk RT > 0.5) to be completely replicated. Indeed, this is not the case, as about 95-99% of early bins have been replicated at this stage (Fig. 6a,b and Supplementary Fig. 5a,b, right panel).

**Figure 6.**
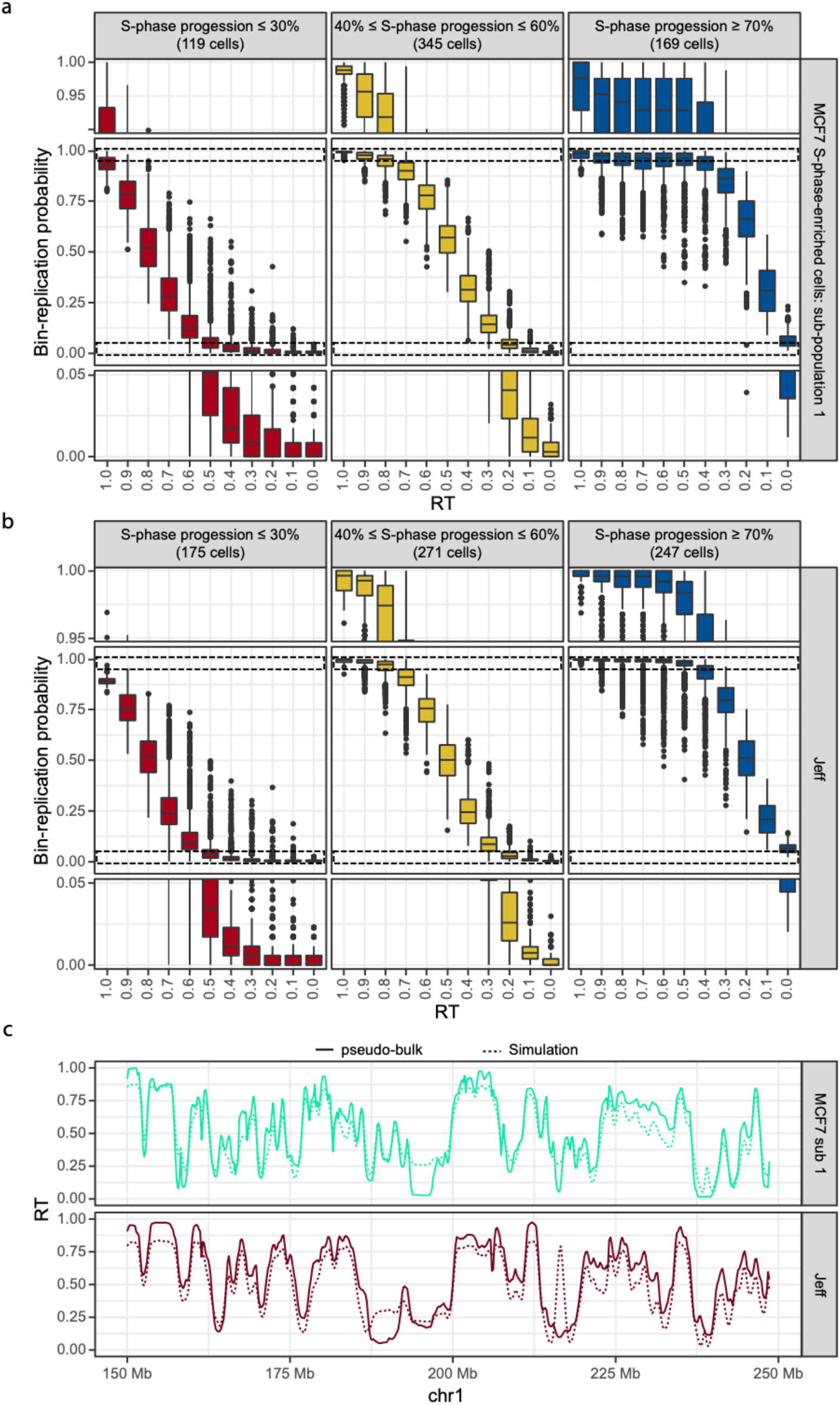
scRT data support a stochastic model of replication. **a-b** Boxplots reporting the replication probability (y-axis) relative to its pseudo-bulk RT (x-axis, 1 is early and 0 is late) for S-phase-enriched cells of MCF7 sub-population 1 (a) and Jeff (b) at different S-phase stages calculated in 200 kb bins. The middle panel of each plot shows the whole distribution, and the top and bottom panels are zoom in for the extremities of the distributions (indicated with the dashed boxes in the middle panel). **c** Comparison between the pseudo-bulk RT (solid line) and simulated RT (dashed line) for the MCF7 sub-population 1 and Jeff cells. The simulation is based on Replicon^29^ and uses the probability of being replicated within early-S-phase cells (completed up to 30% of their genome replication) for each 200 kb bin as input. Similar results were obtained with scRT data of S-phase-enriched MCF7 sub-population 2 and HeLa cells (Supplementary Fig. 5). Spearman correlation in Supplementary Fig. 5d.

Importantly, for a bin, its probability of replication within early-S-phase cells is highly correlated with its population RT (Fig. 6a, b and Supplementary Fig. 5a,b, left panel) for all examined cell types, suggesting that the RT of a given genomic region depends on its firing probability within early-S phase as suggested by the stochastic models. To further test this hypothesis, we selected single cells at the beginning of the S-phase (percentage of replication ≤ 30%) and used the obtained scRT data to calculate the replication probability along the genome. These were then used as an input for Replicon, a stochastic replication simulator^29,30^, to simulate the RT program along the genome (Fig. 6c and Supplementary Fig. 5c). The obtained simulated RT profiles are highly similar (Spearman correlation ≥ 0.857) to the pseudo-bulk RT of the corresponding cell lines (Fig. 6c and Supplementary Fig. 5c,d), demonstrating that the replication signals detected in early-S-phase cells for the late replicating regions are real biological signals instead of technical noise. Our results strongly support the notion of a stochastic RT program.

### Kronos scRT can extract scRT from various single-cell DNA sequencing data

Although, we performed experiments on scWGS, created ad hoc for the scRT analysis, this is not a requirement when using Kronos scRT. As long as the single-cell sequencing data maintain the copy number information and they are derived from cycling cells, Kronos scRT can process them and extract scRT profiles. Amongst the published datasets, scHi-C data generated in Nagano et al. 2017^31^ are a perfect example to demonstrate this. The dataset includes cycling mESC grown either on a feeder layer in ES-DMEM with fetal bovine serum (mESC Serum) or without feeders and adding PD and CHIR inhibitors, two inhibitors that favor the maintenance of a naive ground state for mESC (mESC 2i)^31^. Despite being paired-end sequencing data, due to their nature, they were loaded into Kronos scRT as single-end data. Using Kronos scRT, we identified 312 G1/G2-phase and 329 S-phase cells for the 2i, and 76 G1/G2-phase and 130 S-phase cells for the serum condition. We then calculated the scRT of the S-phase cells and the pseudo-bulk RT profiles (Fig. 7a). The data obtained from scHi-C correlates well with the bulk RT and the pseudo-bulk RT of the mESC analyzed previously from scWGS of sorted mid-S-phase cells (Spearman correlation > 0.847) (Fig. 7a,b and Supplementary Fig. 2a). Furthermore, having more cells well distributed across the S phase allowed us to calculate T_width_ values with the same number of replication categories used for the human samples (Fig. 7c). Our result confirmed that, like human cells, both mESC samples show a tighter replication timing at the beginning and end of the S phase compared to mid-S phase, suggesting that this is an important common feature for mammalian cells.

**Figure 7.**
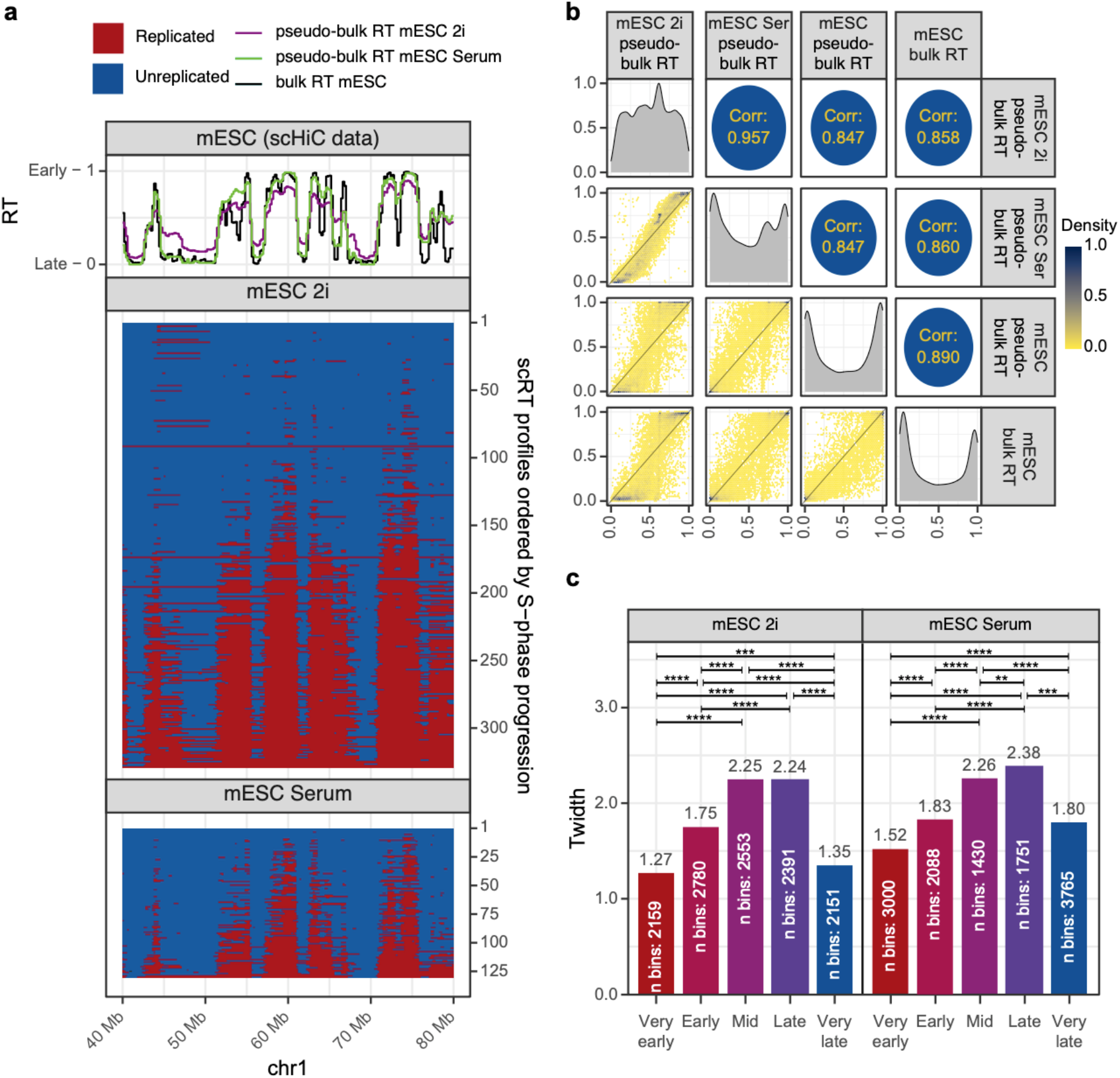
Analysis of scRT from scHi-C data with Kronos scRT. **a** On top: pseudo-bulk RT profile from mESC grown in 2i (purple) or in serum (green) compared to bulk RT (black). In the bottom panel, the scRT profiles ordered from top to bottom by replication percentage of each cell. **b** Pair wise comparisons of mESC pseudo-bulk RT and bulk RT profiles. Same as in Fig. 2b. **c** T_width_ calculated for 5 RT categories based on the corresponding psudo-bulk RT values as in Fig. 3d. P-values were calculated with the Compare TW module as before (* < 0.05, ** < 10^−2^, *** < 10^−3^, **** < 10^−4^).

## Discussion

While RT studies have entered the single-cell era, various limitations have confined the outcoming conclusions to the volume of cells included. Here, we overcome this obstacle of scalability by using a microfluidic-based system to generate new data along with our unified computational workflow (Supplementary Fig. 1a).

We demonstrated how Kronos scRT allows rapid extraction and analysis of scRT from various single-cell DNA sequencing data (such as scWGS, scHi-C) at a 200 kb resolution. Most importantly, Kronos scRT can directly extract scRT with data from asynchronous cycling cells, which avoids time consuming experimental procedures (such as cell sorting into G1 and different periods of S-phase, manual processing on 96 well plates, etc.) as in the currently existing approaches^21–23^. It is to be noted that a similar strategy has been followed by Massey and Koren^32^, who used the 10x Genomics system to study scRT. Our down-sampling analysis indicates that, with Kronos scRT, about 0.75 million reads are sufficient to obtain a suitable scRT at a good resolution for a diploid human genome (Fig. 1d). With the current sequencing cost (∼15 k€ for 10G 100 bp paired-end reads on NovaSeq), we can reach close to a price of ∼1€ per single cell. We successfully applied Kronos scRT to obtain thousands of high-quality scRT profiles from various mouse and human cell types, which allow us to study the stochastics replication events at an unprecedented depth. In agreement with recent ORM data obtained from synchronized early S-phase cells^24^, our data obtained directly from asynchronized cells also support a stochastic replication model. This indicates that the early replication events observed within the late replicating regions detected by ORM are not resulted from the activation of dormant origins due to cell synchronization.

Our analysis demonstrates that we can apply dimensionality reduction to scRT/scCNV profiles in order to identify sub-populations inside heterogeneous samples. By renormalizing the copy-numbers of MCF7 replicated S-phase-cell sub-populations with their corresponding G1/G2-phase counterparts, we have successfully unveiled two different, although relatively similar, RT programs (Fig. 5 and Supplementary Fig. 1d).

Moreover, dimensionality reduction analysis on scRT profiles gives a reconstruction of the replication timeline spanning from early to late S phase of a given population, therefore forming pseudo-trajectories. Deconvoluting cell-to-cell heterogeneity is an important factor to take into consideration, especially for the data from cancer specimens, where normal and mutated cells coexist and the latter group has gone through multiple rounds of random mutation and clonal expansion^33,34^. Furthermore, the possibility of identifying subpopulations in the context of a tissue would prompt the study of the RT program of cells that cannot be cultured *in vitro* and for which specific markers for selection are not available.

Surprisingly, although the two sub-populations of MCF7 cells identified in this study containing significant differences of CNVs (Fig. 4 and Supplementary Fig. 3), their RT profiles are highly correlated (Fig. 5) (R=0.946), suggesting that the RT program is an extremely robust process and different copies of the same chromosome follow similar replication program. This is in agreement with previous reports showing that the variation of RT between homologs (using hybrid musculus, i.e. 139 x Castaneus cells) within the same cell is close to the variation of RT between different cells of the same cell types^21^. Due to the low SNP coverage and low sequencing depth per single cell in our samples, we are not able to calculate the scRT of different homologs. Therefore, at the current technical stage, we can only analyze them as a binary system (i.e. replicated and unreplicated). New algorithms thus need to be developed to obtain the haplotype-resolved scRT in order to explore the variation of replication program between homologous chromosomes in a more general context, such as in normal human diploid cells or cancer cells with complex karyotype. This will help to further investigate the link between structure variation and RT change, and its role in human diseases, such as during cancer developments.

Additional studies are necessary to reveal the molecular mechanisms contributing to the degree of RT stochasticity. The combination of other single-cell omics data, along with scRT, will provide further insights in DNA replication regulation. It should be noted that, RT has long been correlated with chromatin organization. In particular, both RT obtained from population and single cells show that early/late replicating domains are associated with the A/B compartment structure^21,23,35–38^, although the direct causal relationship remains unclear. Interestingly, although changes in RT and compartments during mESC differentiation are tightly linked, they are separable in certain contexts^23^. Comparison of haplotype-resolved bulk RNA-seq and scRT data from mice has indicated that allelic replication asynchrony is frequently but not always associated with allelic expression^22^. Multiple mechanisms might cause extrinsic (cell-to-cell) and intrinsic (homolog-to-homolog) replication variability, and whether the observed association between DNA replication and gene transcription results from the transcription per se or it is indeed due to the active chromatin state of active gene remains an open question^9,39^. Therefore, it’s important to further perform multi-omic studies (simultaneous analysis within the same single-cells) to provide a better understanding of replication kinetics. Potentially, the remarkable simplicity of our Kronos scRT approach, which allows an increased number of cells to be integrated, can also be easily combined with the analysis of single-cell RNA-seq, scHi-C, CpG methylation and chromatin accessibility^40–43^, among others, offering an opportunity to study RT at both a multi-omic scale and the single-cell level. Furthermore, these single-cell multi-omic data could also be used to extract the RT landscapes while simultaneously fulfilling their original purpose (e.g. chromatin accessibility for scHi-C and scATAC, transcription for scRNA-seq, etc). We therefore underline the need to increase the number of such studies in order to better comprehend DNA replication control and its stochasticity at the single-cell level.

## Methods

### Cell culture

Estrogen Receptor (ER)-positive breast cancer MCF7 cells were cultured and treated as mentioned in^44^. In brief, cells were maintained in complete media (DMEM supplemented with 10 % FBS, 50 U/mL penicillin and 50 μg/mL streptomycin) in 5% CO_2_ at 37 °C. Estrogen (E2) treatment was performed after hormone starvation. Briefly, cells were plated at ∼25% confluent in complete media for at least 16 h, rinsed with PBS for 3 times, and then hormone starved for 48 h in hormone-free media (DMEM without phenol red supplemented with 10% charcoal stripped FBS (Dutscher), 50 U/mL penicillin and 50 μg/mL streptomycin) before treated with 100 nM E2 (dissolved in EtOH) for 24 h. Cells were harvested post E2 treatment (70-80% confluence) via trypsin detachment. HeLa S3 cells were cultured in DMEM high glucose media with 10% FBS, while JEFF cells were grown in RPMI 1640 supplemented with 5% FBS. HeLa S3 cells were harvested via trypsin detachment around 70-80% confluent, while JEFF cells were harvested at 0.7-0.8 × 10^6^ cells/mL by centrifuge.

### Single-cell copy number variation sample preparation

Libraries were produced starting from exponentially growing cells. If mentioned, a step of S-phase enrichment through FACS sorting was performed. Cells were processed using the 10x Genomics Chromium single-cell CNV solution accordingly to manufacturer’s instruction. Libraries were sequenced on an Illumina Novaseq 6000 using PE100, aiming to obtain 2 million unique reads per single cell.

### S-phase enrichment through Fluorescence-activated cell sorting

Exponentially growing cells were collected and stained with 20 μg/mL Hoechst 33342 in complete media at 37 °C for 1 h. Stained cells were rinsed twice with PBS, clumps were removed by passing each sample twice through a 40 μm strainer. The resulting single cell suspension was stained with 50 μg/mL PI before FACS. PI-positive cells were discarded, and cell-cycle stages were estimated according to the Hoechst signal. Gates were positioned in order to collect S-phase cells and partially G1- and G2/M-phase cells. Sorted cells were collected in a 15 mL tube with 1 mL complete culture media, rinsed once in PBS (Ca^2+^ free, Mg^2+^ free)-0.04% BSA and then run on a 10x Genomics Chromium with single-cell CNV solution kit as previously described.

### Fluorescence in situ Hybridization (FISH)

For FISH analysis, cells were treated with colcemid (100 ng/ml, Roche) for 3 h and mitotic cells were collected by mitotic shake-off after a short trypsin treatment and centrifuged at 1000 rpm for 10 min. Cell pellets were resuspended in 75 mM KCl and incubated for 15 min in a 37°C waterbath. Carnoy fixative solution (methanol/acetic acid, 3:1) was prepared and 1:10 volume added on the cells, before centrifugation at 1000 rpm for 15 min. Cells were then fixed for 30 min at room temperature in the carnoy solution, centrifuged and washed once more with fixative. Minimum volume of fixative was left to resuspend the pellet and cells were dropped onto clean glass slides. FISH staining was performed following manufacturer’s instructions (MetaSystems) using chromosome painting and centromere enumeration probes to specifically identify the chromosome 3 (Metasystems probes). The Metafer imaging platform (MetaSystems) was used for automated acquisition of the chromosome spread. Picture triplets were merged with Fiji (v2.1.0) and the resulting images were manually scrutinized for chromosome 3 enumeration. Representative pictures were acquired using a Deltavision Core system (Applied Precision).

### 10x Genomics data processing

Data produced with 10x Genomics system (various human cell lines) were processed using Cell Ranger DNA (10x Genomics software, version 1.0.0). The resultant bam file was subset using 10x Genomics subset-bam version 1.0 to obtain single cell bam files. Duplicated reads were removed using Picard MarkDuplicates (version 2.6.0, http://broadinstitute.github.io/picard) and the resulting files were used as input for the CNV module of Kronos scRT (Supplementary Fig. 1a).

### Trimming and aligning reads

The fastqtoBAM module of Kronos scRT uses demultiplexed fastq files as input and removes standard adaptors from reads. Adaptor trimming is performed using Trim Galore (version 0.4.4, https://www.bioinformatics.babraham.ac.uk/projects/trim_galore/), a modified version of cutadapt and FastQC. After trimming, reads are aligned to the provided reference genome (in our study, human genome version hg38 and mouse genome version mm10 were used) using RBowtie2 package (version 1.4.0) reporting only the best mapping for each read. SAM files are then sorted, converted into BAM files using Rsamtools (version 1.34.1) and deduplicated using Picard MarkDuplicates (version 2.6.0). The fastqtoBam module was used to process all the mouse data analyzed in the current study.

### Calculate bin mappability and GC content for copy-number estimation

The binning module of Kronos scRT is used to calculate mappability and GC content in each genomic bin. This information is later used by the CNV module to normalize read counts and select which bins will be considered for further analysis. By default, bin size is 20 kb, but, depending on the sample average sequencing dept, it can be adjusted by the user. Moreover, only autosomal chromosomes are used by default, but the user can decide to keep either or both sex chromosomes as well. GC content is simply calculated as the frequency of C and G in the reference sequence belonging to a bin.

To calculate mappability, this module simulates 1X coverage reads from a reference genome, adds mutations with an error rate of 0.1% (that can be adjusted by the user to fit the error rate of their own datasets^45^) and re-maps the reads on the reference genome using Rbowtie2 with the same settings used in the fastqtoBAM module. Read parameters such as length of the reads, single-end or paired-end reads and insert size are estimated from the BAM files of the single cell experiment or they can be manually set by the user. The mappability of the bin n (*M*_*n*_) is therefore calculated as the number of remapped reads of this bin (*Rr*_*n*_), divided by the number of reads that were originally generated from the same location (*Rs*_*n*_) (i).

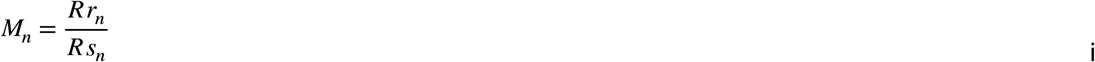

### CNV calling and intracellular bin-to-bin variability

The CNV module of Kronos scRT counts the number of high-quality reads, mapping quality score ≥ 30, over the bins generated by the binning module: for CNV calculation of pair-end reads, if reads of a pair mapped in the same bin, they are counted only once, otherwise, they are counted independently. Cells with less than 2*10^5^ reads are discarded (the user can manually adjust this threshold). Regions with a mappability lower than 0.8 or higher than 1.5 are excluded from the rest of the analysis (the user can as well provide a list of blacklisted genomic regions and/or change mappability thresholds). Read counts are then adjusted based on mappability as in formula (ii).

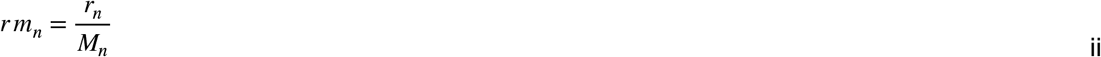

Where *r*_n_ is the read count on the bin n, *M*_*n*_ is the mappability of the bin *n* obtained from (i) and *rm*_*n*_ is the adjusted read count based on its mappability.

Read counts is then corrected for the GC content bias (formula iii).

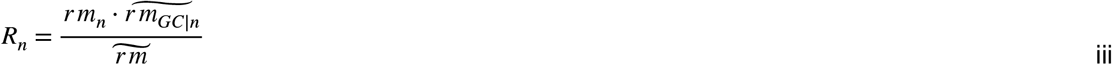

Where *R*_*n*_ is the normalized read count of the bin n, *rm*_*n*_ is the adjusted read counts (ii), *GC*|*n* represents all the bins with the same GC content of the bin n and the tilde represents the median. The CNV module will first calculate the bin-to-bin variability. To do so, it will re-bin the genome into 500 kb bins and calculated the total read count for each bin. Then it will proceed calculating the Depth Independent Median Absolute deviation of Pairwise Differences (DIMAPD) as defined in 10x Genomics CNV solution. Assuming that the majority of analyzed cells is in G1/G2 phase, in which the bin-to-bin variation is minimal, DIMAPD values are fitted to a gaussian distribution and cells that are statistically different with high DIMAPD (formula iv, v, vi and vii) are therefore considered belonging to S-phase.

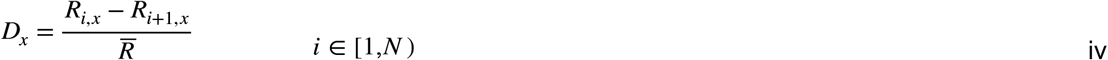

Where *D*_*x*_ is a vector conting differences between neighburing 500 kb bins for the cell *x, R*_*x*_ is a vector containing the number of reads in 500 kb bins for the cell *x*, and the index *i* identifies a bin (ranges betwen 1 to the total number of bins -1).

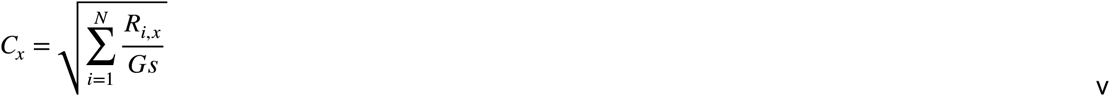

Were *C*_x_ is the square root of the coverage, *N* is the number of 500 kb bins, *R*_*i,x*_ is the number of reads in the bin *i* in the cell *x* and *Gs* is the genome size in Mb.

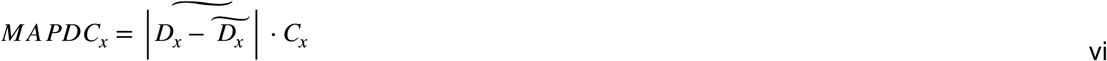

Where *D*_*x*_ comes from (iv), and *C*_*x*_ from (v).

The MAPDC increases linearly with the square root of the cell coverage, therefore we normalize it as follow to obtain the DIMAPD:

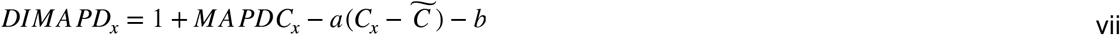

Where *MAPDC*_*x*_ is defined in formula (vi), *C*_*x*_ in formula (v) and *C* is a vector containing the values of *C*_*x*_ for all the cells in the experiment. *a* and *b* are two coefficients estimated through linear fitting of MAPDC in function of the cell coverage distance from the median coverage of the experiment. CN are called starting from 20 kb bin tracks that are smoothed and segmented using circular binary segmentation (CBS) algorithm from the R package DNAcopy (version 1.56.0). CN is then estimated through the minimization of the following target function as suggested by 10x Genomics (formula viii).

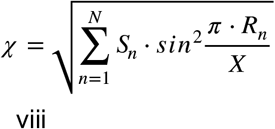

Where *S*_*n*_ represents the size of the segment *n, R*_*n*_ represents the read count of segment *n*, and *X* is a number between the 5^th^ and the 95^th^ percentile of the read counts of all the segments in a cell. Each local minimum of this equation is a possible solution to calculate CN. A filter on local minima that lead to unreasonable mean ploidy (formula x) is therefore applied and CN is calculated (formula ix). The filters used in this study can be found in Supplementary Table 2 (Ploidy limits). For the sorted G1 and Mid-S cells of the mouse datasets the closest ploidy to 2 was selected.

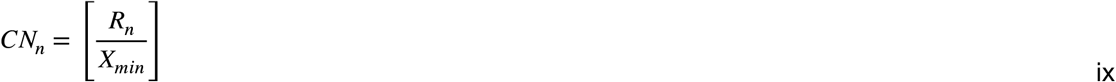

Where *X*_*min*_ is the value of for *x* which 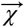 is minimized (formula viii), *R*_*n*_ is the read count in the segment *n*, and *CN*_*n*_ is an integer that represent the copy number of the segment *n*. The mean ploidy of a cell can then be calculated as follow:

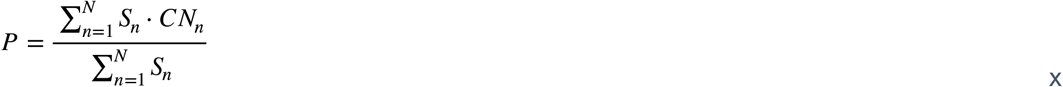

Where *P* is the mean ploidy, *S*_*n*_ is the size of the segment *n*, and *CN*_*n*_ is the copy number of the segment *n*. The difference between the absolute minima and its closest relative minimum is used to evaluate how good is the CN calling, for values below 2 the CN is not considered reliable (ploidy confidence). Negative values of ploidy confidence are imposed as suggested by 10X Genomics. Bins included in the provided blacklist are removed, as done here with the binning for the mESC and mNE-7d cells.

### Single-cell replication profiling and scRT calculation

As already mentioned, the DIMAPD parameter can be used to distinguish the replicating cells (i.e. S-phase cells) from those which are not (i.e. G1/G2-phase cells). The automatic threshold is a reasonable choice if the cell population has not been sorted to enrich for the S phase. For the S-phase enriched cells, the diagnostic module can be used to select more adequate thresholds manually. The thresholds used in this study are reported in Supplementary Table 2. If available, FACS metadata can be integrated through the WhoIsWho module of Kronos scRT.

As shown in Fig. 1b, the function that we use to identify CN (formula viii) introduces some constraints in the calculation of mean ploidy. Firstly, it is not possible to distinguish between G1 and G2 cells that co-occupy the same area (Fig. 1b, blue population). Secondly, the S-phase cells are split into two (Fig. 1b, green population): with the first part that progresses normally, while the second part is approaching the G1/G2 population from the left side of the plot as indicated by the two arrows. Therefore, Kronos scRT diagnostic module calculates two parameters to correct S-phase populations. Preferentially, the program tries to reunite the S phase in a monomodal distribution in which the ploidy variability is maximized. When this is not possible, parameters are chosen in order to create a bimodal distribution with a minimized ploidy variability. The user can manually set these parameters.

The CN of each segment is as well corrected based on these values. Accordingly to our down-sampling (Fig. 1d and Supplementary Fig. 1b,c), cells with low coverage were filtered out. Coverage thresholds for each dataset are reported in Supplementary Table 2.

The genome is then binned again, in this case, in bins of 200 kb to calculate scRT, but the size should be adjusted based on sequencing depth. A weighted median CN is then calculated, where the weights are the sizes of overlap between each 200 kb bin and the previously calculated segments.

The G1/G2-phase population is used to calculate a median pseudo-bulk CN profile and it is used to normalize each cell in S phase as follow (formula xi):

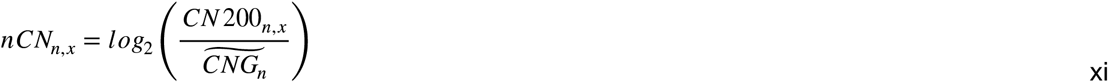

Where *nCN*_*n,x*_ is the normalized copy number of the bin *n* in the S-phase cell *x. CN200*_*n,x*_ is the copy number of the bin *n* in the S-phase cell *x* before normalization, and *CNG*_*n*_ is the CN of the bin *n* in all the G1/G2-phase cells.

Each S-phase cell profile is then binarized. To do so, Kronos scRT identifies a *nCN* value for which the following target function is minimized (xii) (Fig. 1e).

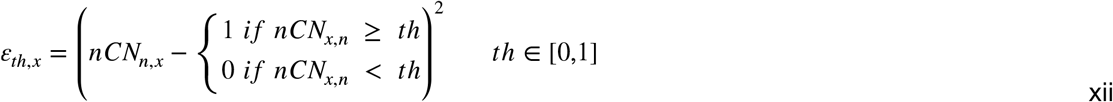

Where *ε*_*th*_,_*x*_ is the Euclidian distance using the threshold *th* for the cell *x* and *nCN*_*n,x*_ is obtained from (xi). Once the threshold that minimizes *ε*_*x*_ is identify, we can calculate scRT profiles as follow (xiii).

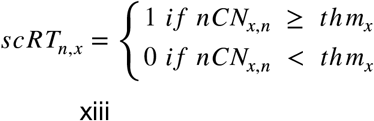

Where *scRT*_*n,x*_ is a binary value representing whether the bin *n* in the cell *x* has been replicated (1) or not (0), *nCN*_*n,x*_ comes from (xi) and *thm*_*x*_ is the *th* for each *ε*_*x*_ is minimized in the cell *x*. Simple matching coefficient distances are then calculated for each pair of cells. The population is filtered to remove cells that diverge by at least 25% from 60% of the single-cell population (Fig. 1f). Cells are then sorted by percentage of replication and tracks are averaged within each bin of replication percentage. In order to ensure a symmetrical distribution, excess cells (i.e. very early and/or very late) are filtered out. Replication tracks per percentage interval are then averaged together to create pseudo-bulk RT and compared with bulk RT. In this study, bulk RT of mESCs and NE-7d issued from BrdU-IP samples had coordinates converted to mm10 with R package liftOver (v1.10). Human bulk RT were converted from hg19 to hg38 using ucsc-liftOver tool (v366).

### Studying variability and sample differences

To study cell-to-cell variability, Kronos RT calculates T_width_: the time needed for genomic regions to be replicated from 25% to 75% of cells in a S-phase lasting 10 h. The module Compare TW of Kronos RT allows user to apply a null hypothesis test though bootstrapping with H_0_: T_width_group1_ = T_width_group2_ and with H_1_: T_width_group1_ ≠ T_width_group2_. To do so, it randomly assigns the bins belonging to two groups to either of them, keeping the total original number of bins in each group constant. Newly assigned bins are then used to calculate the absolute difference between T_width_group1_ and T_width_group2_ and then compare it with the real difference (xiv).

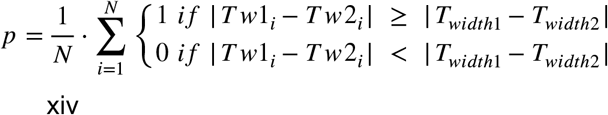

Where *p* is the p value, *N* is the number of iterations, by default 10^4^, *Tw1*_*i*_ and *Tw2*_*i*_ are the T_width_ calculated, respectively, for the two groups in the iteration *i*, while *T*_*width1*_ and *T*_*width2*_ are the values of the real groups.

### Down-sampling

To test the CN calling stability in function of the dept of sequencing, we selected G1/G2- and S-phase cells from each experimental setting: i.e. 10x Genomics system, scWGA and scHiC. Down-sampling was performed using Picard DownsampleSam (version 2.6.0, http://broadinstitute.github.io/picard). For each down-sampling coverage, cells with higher RPMb were used. RPMb thresholds were set in our study as the value at which at least 75% of cells have a ploidy estimation that does not differs more than 5% from the original value. If the users do not define a specific threshold, Kronos scRT applies a threshold of 160 RPMb per haploid genome by default, which approaches the highest threshold we found from different datasets analyzed in the current study (Fig. 1d and Supplementary Fig. 1c).

### Dimension Reduction

Kronos DRed is the dimension reduction module, which uses genome-wide scCNV or scRT data to transform the data and provide a low-dimensional representation reflecting the important features (e.g. cell type, cell population, etc.). For CNV data, original values are used, while for RT data, simple matching coefficient distances calculated from R package ade4 (v1.7) are used. t-SNE^46^ and UMAP^26,27^ are performed with R packages Rtsne (v0.15) and umap (v0.2.7), respectively. For t-SNE, perplexity corresponds to a fiftieth of the number of cells or a minimum value of 10 accordingly, theta to 0.25, and partial PCA is performed allowing t-SNE coordinates to be calculated under 5,000 iterations.

### MCF7 sub-population separation

Kronos RT (option --extract_G1_G2_cells) was used to generate complete S and G1/G2 phase CNV in 200 kb bins. Bins containing missing values or those belonging to sex chromosomes were removed. Dimension reduction of the resulting data was performed under UMAP using R package umap (v0.2.7, option random_state=20210813). 2D UMAP coordinates were projected along with the percentage of replication of the cells for G1/G2 and S-phase distinction. Cells were labelled based on manually attributed cut-off coordinates from the UMAP projection. For each resulting group, genome-wide scCNV data was visualized (Fig. 4b and Supplementary Fig. 3d), which allowed manual attribution of S-phase groups to their corresponding G1/G2 groups and thus, correct normalization of these groups in the downstream scRT analysis.

### Replication Timing Simulation

We used the Replicon simulation code^29^ to simulate the replication timing profiles. The Replicon simulator uses the initiation probability landscape (IPLS), i.e. the relative probability of initiating at any point in the genome, as input. In our simulations, we used the probability of being replicated within early-S-phase cells (completed up to 30% of their genome replication) for each 200 kb based on the scRT data of the corresponding cell types. We used the same setting for other parameters as in our previous publication^24^ followed the suggestion of the original paper^30^.

## Data availability

The scWGS data of mouse were obtained from GSE108556. The scHi-C data of mouse were obtained from GSE94489. The bulk RT data were obtained from GSM923442 for MCF7, GSM923449 for HeLa, GSM923451 for GM12878 (B-lymphoblastoid cell line) and GSE108556 for mESC and NE-7d cells. The mm10 blacklist can be located at https://github.com/Boyle-Lab/Blacklist. The raw and processing generated in the current study will be submitted to SRA and GEO, respectively.

## Code availability

Kronos scRT is available on github (https://github.com/CL-CHEN-Lab/Kronos_scRT).

### Acknowledgements

Work of C.L. C lab is supported by the YPI program of I. Curie, the ATIP-Avenir program from Centre national de la recherche scientifique (CNRS) and Plan Cancer from INSERM, the CNRS 80|Prime interdisciplinary program, the Agence Nationale pour la Recherche (ANR) and Institut National du Cancer (INCa). J.M.J. is supported by a PSL-Qlife fellowship (ANR-17-CONV-0005). High-throughput sequencing was performed by the ICGex NGS platform of the Institut Curie supported by the grants ANR-10-EQPX-03 (Equipex) and ANR-10-INBS-09-08 (France Génomique Consortium) from the Agence Nationale de la Recherche (“Investissements d’Avenir” program), by the ITMO-Cancer Aviesan (Plan Cancer III) and by the SiRIC-Curie program (SiRIC Grant INCa-DGOS-465 and INCa-DGOS-Inserm_12554). The authors would like to acknowledge Geneviève Almouzni for the MCF7 cells, the cell sorting facility of I. Curie for assistance with cell sorting and Dominika Foretek, Marc Gabriel and Ugo Szachnowski for stimulating discussions.

## Author contributions

CLC conceived and planned the study. XW conducted the single-cell experiments with the assistance of MB and LGB. MB and LGB performed the sequencing under the supervision of SB. MS conducted the experiments and prepared the samples for FISH experiment. MD conducted the FISH experiment under the supervision of DF. SG developed the program and JMJ contributed to the development. SG and JMJ performed the bioinformatics analyses. DS performed the RT simulation. CLC supervised the experiments and bioinformatics analyses. SG, JMJ and CLC wrote the manuscript, and all the authors reviewed it.

## Competing interests

The authors declare no competing interests.

## Notes

https://github.com/CL-CHEN-Lab/Kronos_scRT

### Competing Interest Statement

The authors have declared no competing interest.

### Summary of Updates

Detailed benchmarking has been added, which allowed us to reveal the minimum numbers of reads per single cell necessary to build accurate scRT profiles for each analyzed dataset. All figures and tables have been updated.

