## Supplementary Data for "Kronos scRT: a uniform framework for single-cell replication timing analysis"

**Gnan et al.**

### SUPPLEMENTARY INFORMATION

#### Supplementary Table 1. Summary of scRT data analyzed in the current study.

| Organism | Cell lines | Cell enrichment | Method | Number of cells | | | | Ref (raw data) |
|---|---|---|---|---|---|---|---|---|
| | | | | All | Passed filters* | G1/G2 | S-phase | |
| Mouse | mESC | G1 sorted & Mid-S sorted | scWGA | 67 | 67 | 13 | 54 | Ref[22] |
| | mNE-7d | G1 sorted & Mid-S sorted | scWGA | 45 | 45 | 3 | 42 | Ref[22] |
| | mESC 2i | Cycling cells | scHi-C | 1440 | 641 | 312 | 329 | Ref[31] |
| | mESC Serum | Cycling cells | scHi-C | 437 | 206 | 76 | 130 | Ref[31] |
| Human | MCF7 | Cycling cells | 10x scCNV | 447 | 368 | 286 | 82 | This study |
| | MCF7 | S-phase enriched | 10x scCNV | 2321 | 1777 | 424 | 1353 | This study |
| | HeLa | S-phase enriched | 10x scCNV | 752 | 514 | 255 | 259 | This study |
| | Jeff | S-phase enriched | 10x scCNV | 1455 | 1106 | 146 | 960 | This study |

*Based on the simulation results shown in Fig. 1d and Supplementary Fig. 1b,c, stringent thresholds (Supplementary Table 2) were used in our current study to select the high-quality cells, in order to avoid that technical noise be translated as observed cell-to-cell variations.

### Supplementary Table 2. Summary of the parameters used in the analysis.

| Cell | run | Reads type | Ploidy limits | G1/G2-phase threshold | S-phase threshold | Adj. 1st part S-phase | Adj. 2nd part S-phase | Adj. Type | Reads per Mb threshold |
|---|---|---|---|---|---|---|---|---|---|
| HeLa | 1 | PE | [2,8] | 0.90 | 0.85 | 0.950 | 0.540 | Auto | 117 |
| | 2 | PE | [2,8] | 0.90 | 0.60 | 0.950 | 0.537 | Auto | 117 |
| Jeff | 1 | PE | [1.3, 4.3] | 0.80 | 0.75 | 0.950 | 0.550 | Auto | 117 |
| | 2 | PE | [1.3, 4.3] | 0.60 | 0.50 | 0.951 | 0.524 | Auto | 117 |
| MCF7 unsorted | single | PE | [2.5,8] | Automatic | | 0.950 | 0.550 | Auto | 117 |
| MCF7 S-phase enriched | single | PE | [2.5,8] | 0.70 | 0.75 | 0.950 | 0.550 | Auto | 117 |
| mESC 2i | -- | SE | [1.3, 4.3] | 1.00 | 0.90 | 0.950 | 0.550 | Auto | 138 |
| mESC Serum | -- | SE | [1.3, 4.3] | 1.00 | 0.90 | 1.000 | 0.500 | Auto | 138 |
| mESC scRT | -- | SE | ~2 | WhoIsWho | | 0.900 | 0.550 | Manual | 163 |
| NE-7d | -- | SE | ~2 | WhoIsWho | | 0.900 | 0.550 | Manual | 163 |

a

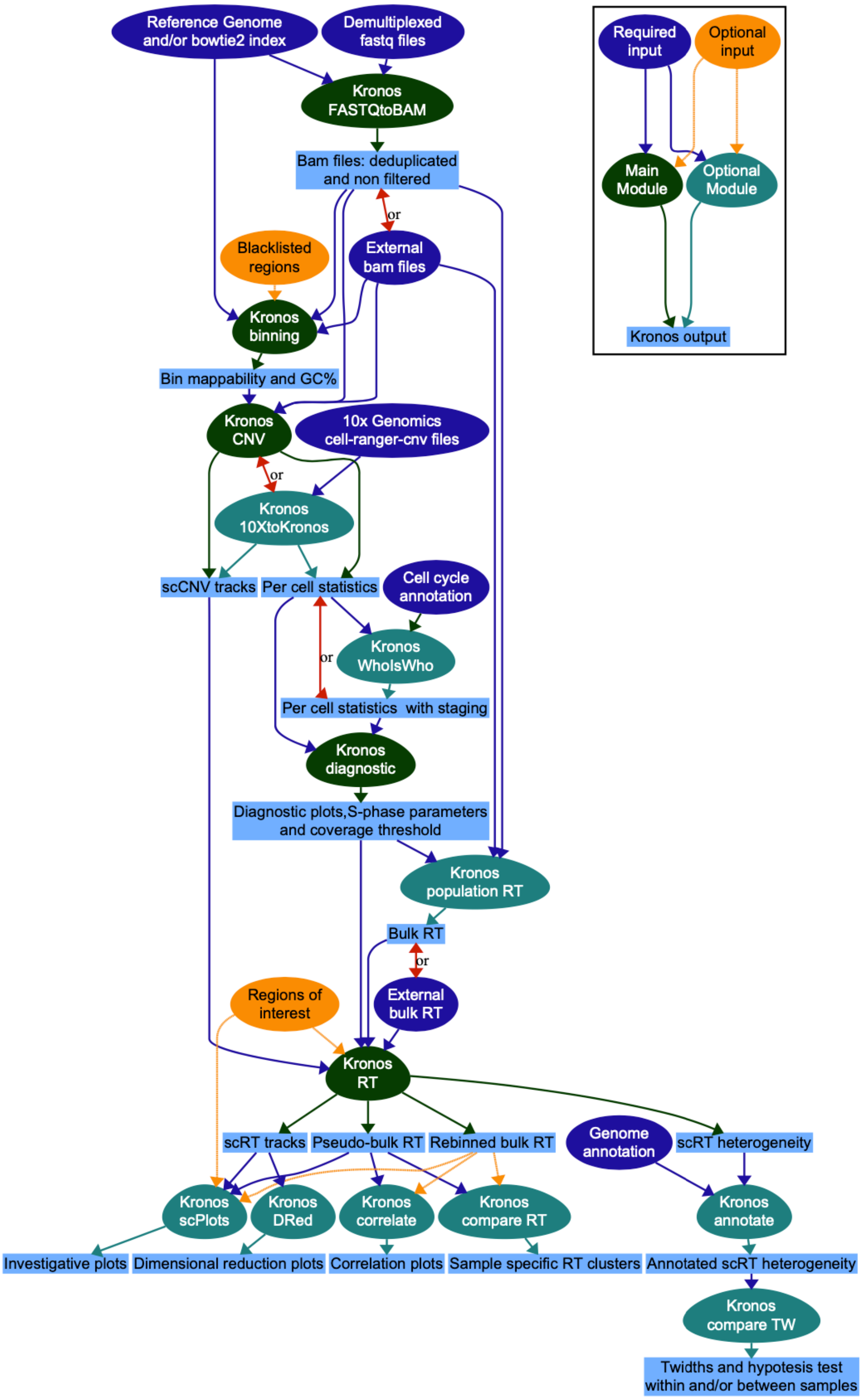

Reference Genome and/or bowtie2 index
Demultiplexed fastq files
Kronos FASTQtoBAM
Bam files: deduplicated and non filtered
or
Blacklisted regions
External bam files
Kronos binning
Bin mappability and GC%
Kronos CNV
10x Genomics cell-ranger-cnv files
or
Kronos 10XtoKronos
scCNV tracks
Per cell statistics
Cell cycle annotation
Kronos WhoIsWho
or
Per cell statistics with staging
Kronos diagnostic
Diagnostic plots,S-phase parameters and coverage threshold
Kronos population RT
Bulk RT
or
Regions of interest
External bulk RT
Kronos RT
scRT tracks
Pseudo-bulk RT
Rebinned bulk RT
Genome annotation
scRT heterogeneity
Kronos scPlots
Kronos DRed
Kronos correlate
Kronos compare RT
Kronos annotate
Investigative plots
Dimensional reduction plots
Correlation plots
Sample specific RT clusters
Annotated scRT heterogeneity
Kronos compare TW
Twidths and hypotesis test within and/or between samples
Required input
Optional input
Main Module
Optional Module
Kronos output

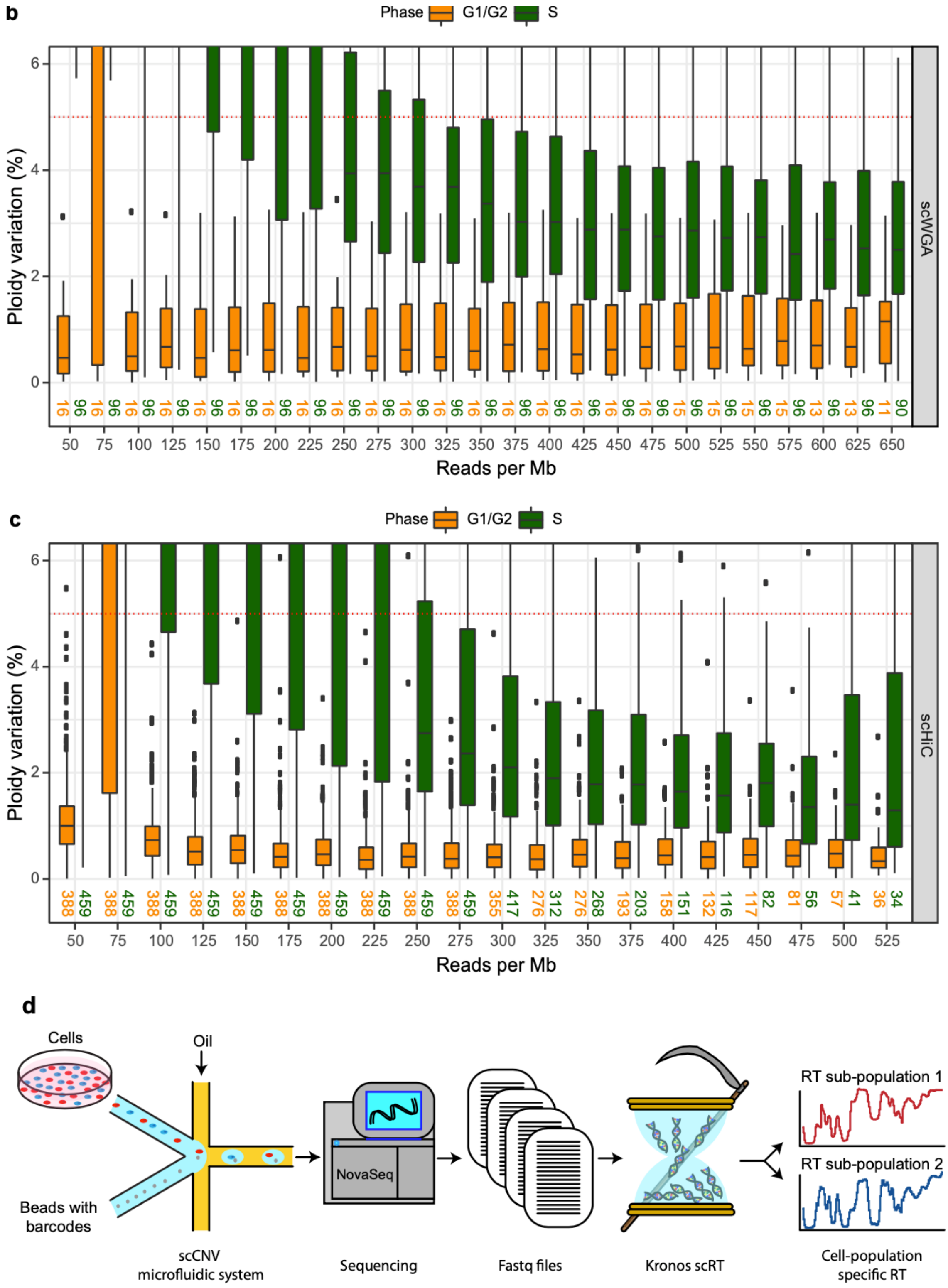


**Supplementary Figure 1. Kronos scRT framework. a** The full pipeline of Kronos scRT with all the developed modules. The input files, the main modules and the optional modules are shown in green, dark blue and light blue, respectively. **b-c** Reads down-sampling as in Fig. 1d for scWGA and scHiC data obtained from Takahashi et al 2019[22] and Nagano et al. 2017[31], respectively. **d** A schematic representation from the experimental set-up to the final data of a mixed population.

a

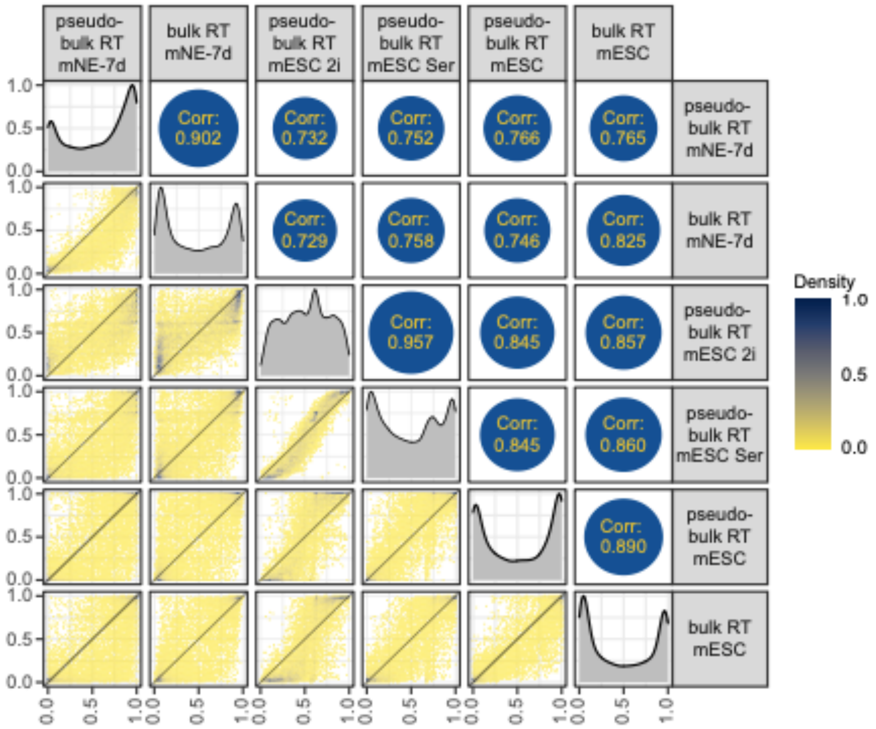
pseudo-bulk RT mNE-7d
bulk RT mNE-7d
pseudo-bulk RT mESC 2i
pseudo-bulk RT mESC Ser
pseudo-bulk RT mESC
bulk RT mESC
Corr: 0.902
Corr: 0.732
Corr: 0.752
Corr: 0.766
Corr: 0.765
Corr: 0.729
Corr: 0.758
Corr: 0.746
Corr: 0.825
Corr: 0.957
Corr: 0.845
Corr: 0.857
Corr: 0.845
Corr: 0.860
Corr: 0.890
Density
1.0
0.5
0.0

b

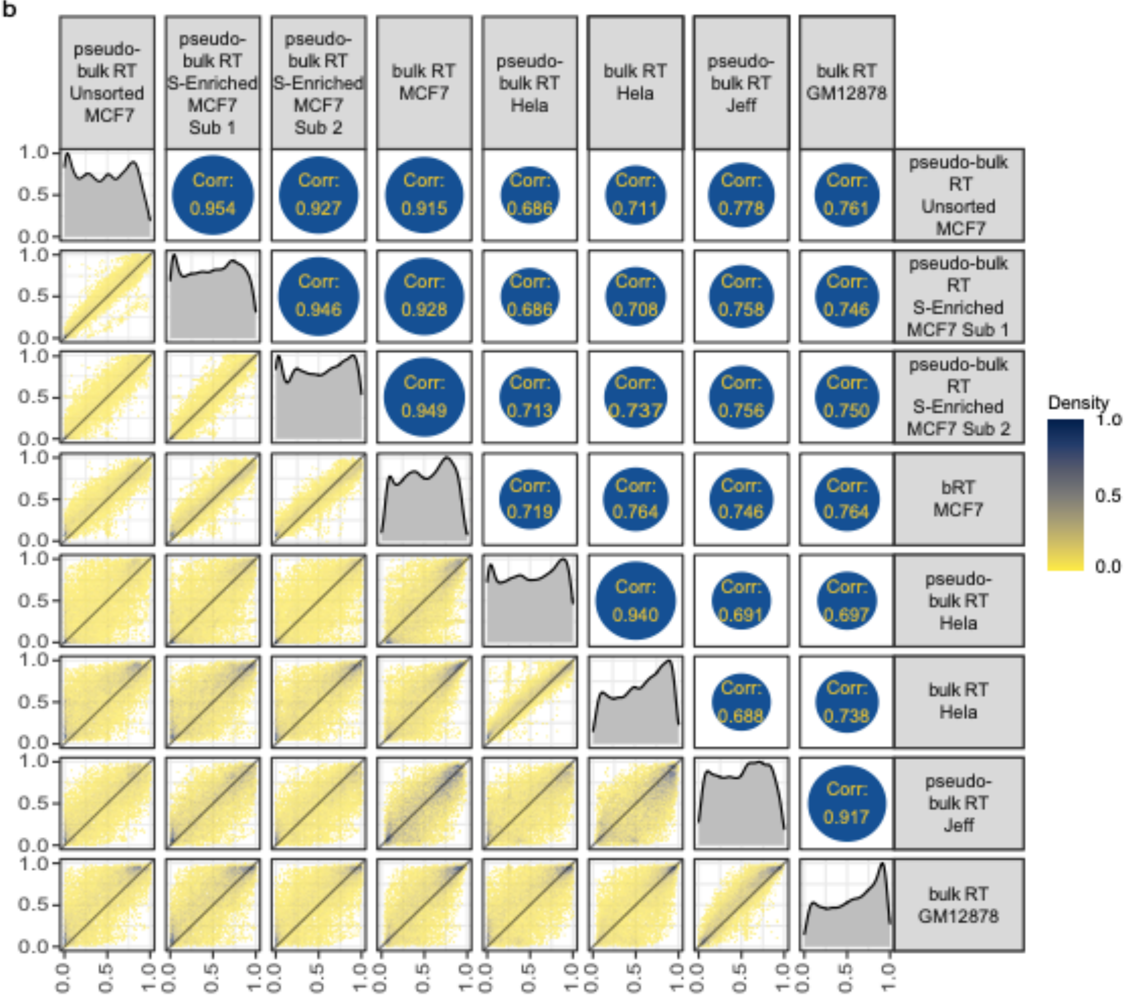
pseudo-bulk RT Unsorted MCF7
pseudo-bulk RT S-Enriched MCF7 Sub 1
pseudo-bulk RT S-Enriched MCF7 Sub 2
bulk RT MCF7
pseudo-bulk RT Hela
bulk RT Hela
pseudo-bulk RT Jeff
bulk RT GM12878
bRT MCF7
Corr: 0.954
Corr: 0.927
Corr: 0.915
Corr: 0.686
Corr: 0.711
Corr: 0.778
Corr: 0.761
Corr: 0.946
Corr: 0.928
Corr: 0.686
Corr: 0.708
Corr: 0.758
Corr: 0.746
Corr: 0.949
Corr: 0.713
Corr: 0.737
Corr: 0.756
Corr: 0.750
Corr: 0.719
Corr: 0.764
Corr: 0.746
Corr: 0.764
Corr: 0.940
Corr: 0.691
Corr: 0.697
Corr: 0.688
Corr: 0.738
Corr: 0.917
Density
1.0
0.5
0.0

**Supplementary Figure 2. Bulk and pseudo-bulk RT comparison. a-b** The upper diagonal of each panel presents a pair-wise Spearman correlation of mouse (a) and human (b) bulk and pseudo-bulk RT profiles. The diagonals present the RT distribution of each sample, while the lower triangles show 2D density plots reporting pair-wise comparisons between samples. Density color code is reported with the heatmap on the right.

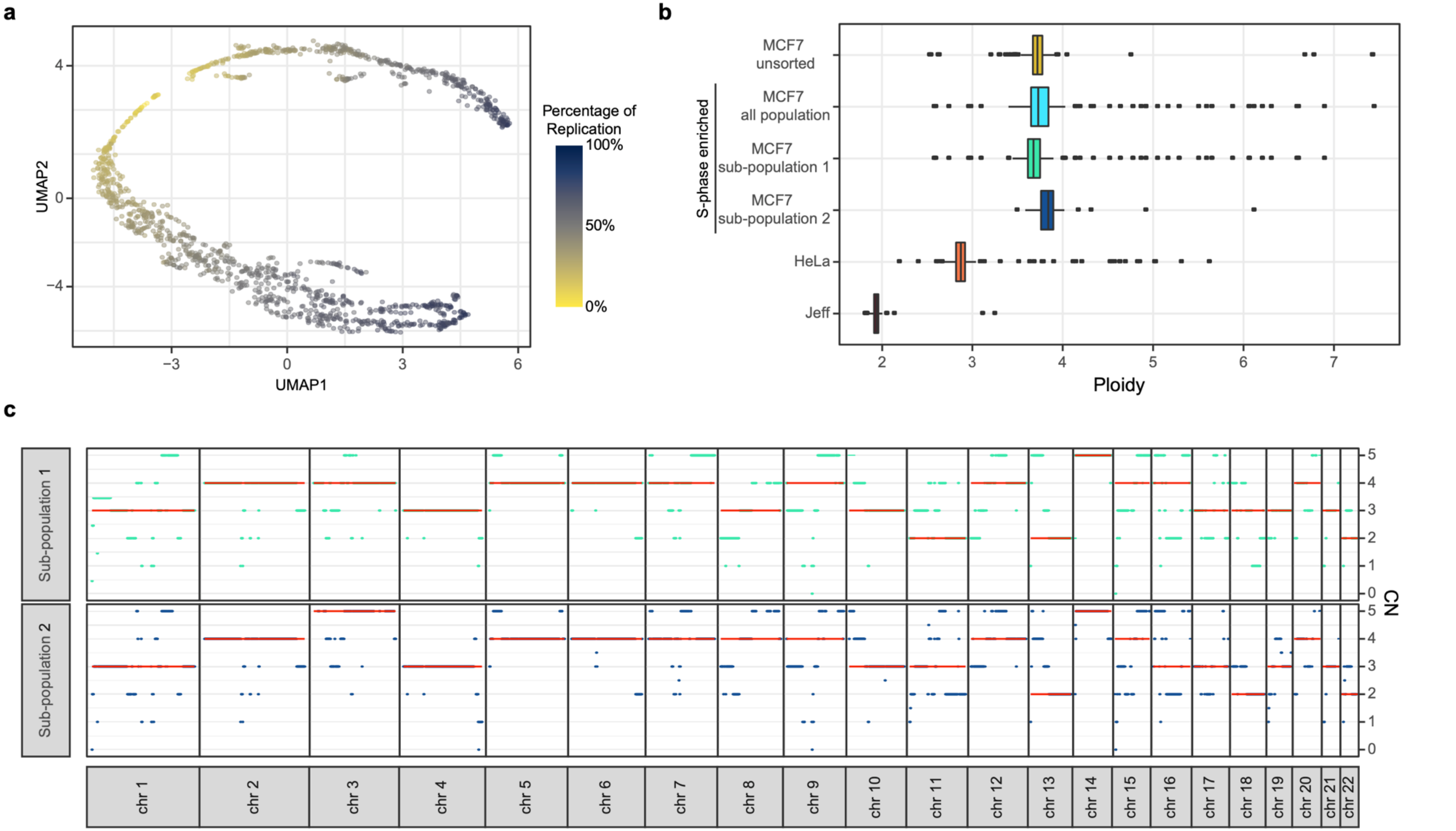

a
b
c
UMAP1
UMAP2
Percentage of Replication
100%
50%
0%
MCF7 unsorted
MCF7 all population
MCF7 sub-population 1
MCF7 sub-population 2
S-phase enriched
HeLa
Jeff
Ploidy
Sub-population 1
Sub-population 2
CN
chr 1
chr 2
chr 3
chr 4
chr 5
chr 6
chr 7
chr 8
chr 9
chr 10
chr 11
chr 12
chr 13
chr 14
chr 15
chr 16
chr 17
chr 18
chr 19
chr 20
chr 21
chr 22

d

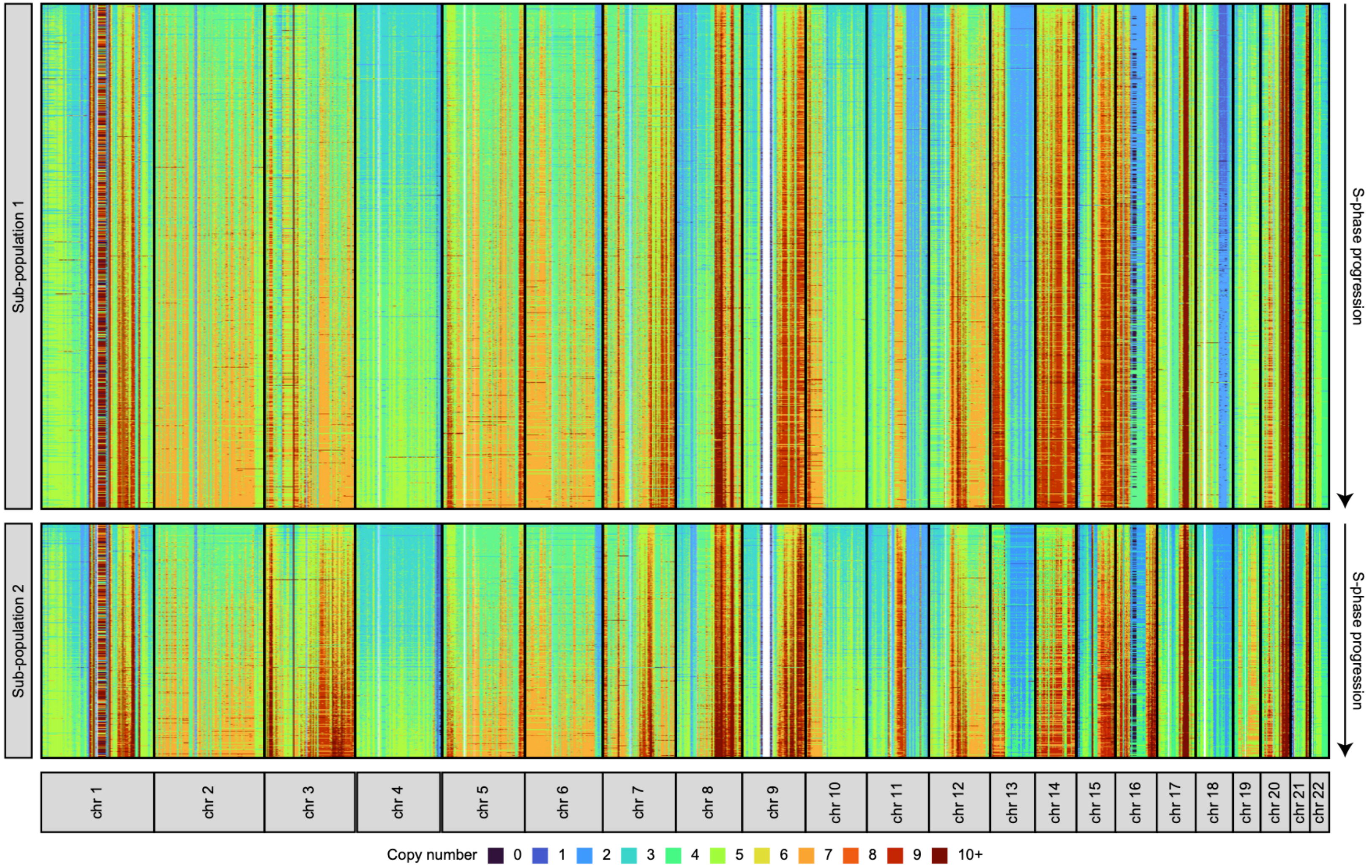
Sub-population 1
Sub-population 2
S-phase progression
S-phase progression
chr 1
chr 2
chr 3
chr 4
chr 5
chr 6
chr 7
chr 8
chr 9
chr 10
chr 11
chr 12
chr 13
chr 14
chr 15
chr 16
chr 17
chr 18
chr 19
chr 20
chr 21
chr 22
Copy number 0 1 2 3 4 5 6 7 8 9 10+

**Supplementary Figure 3. MCF7 cell culture contains two sub-populations. a** Dimensionality reduction analysis of scRT profiles. Each dot represents a single cell and it is color coded based on its percentage of replication. **b** Boxplots reporting mean ploidy of cells calculated by Kronos using the G1/G2-phase population. **c** Median copy-numbers along autosomal chromosomes in G1/G2-phase MCF7 sub-population 1 (aqua) and sub-population 2 (blue) (bin size = 1 Mb). Red lines represent the median CN of the whole chromosome. **d** Copy-numbers along autosomes detected in S-phase MCF7 cells. Cells have been grouped based on the corresponding G1/G2-phase showed in Fig. 4a. Cells were sorted based on S-phase progression.

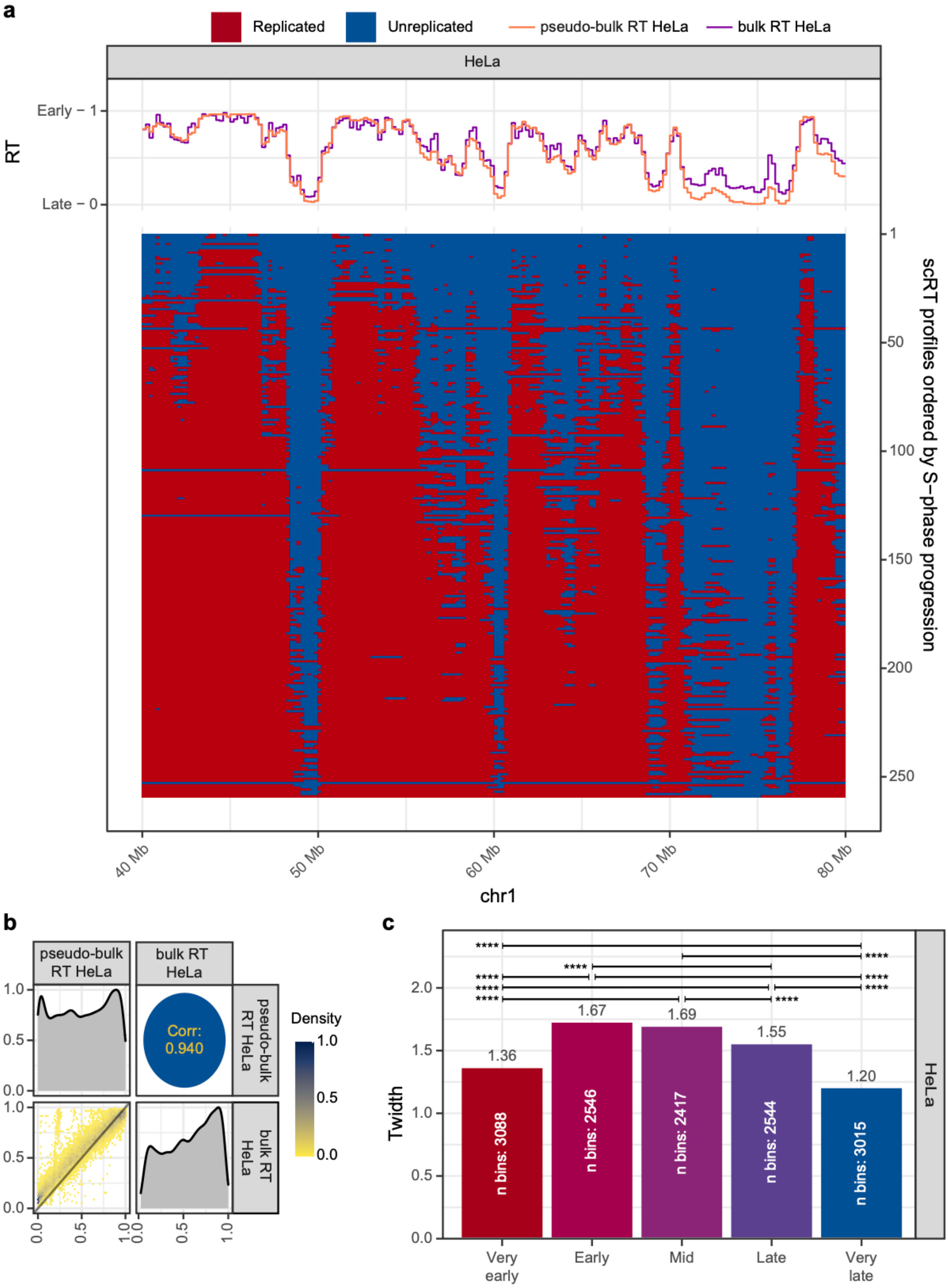
a
Replicated
Unreplicated
pseudo-bulk RT HeLa
bulk RT HeLa
HeLa
RT
Early – 1
Late – 0
scRT profiles ordered by S–phase progression
1
50
100
150
200
250
40 Mb
50 Mb
60 Mb
70 Mb
80 Mb
chr1
b
pseudo-bulk RT HeLa
bulk RT HeLa
Corr: 0.940
Density
1.0
0.5
0.0
c
Twidth
1.36
1.67
1.69
1.55
1.20
n bins: 3088
n bins: 2546
n bins: 2417
n bins: 2544
n bins: 3015
Very early
Early
Mid
Late
Very late
HeLa
****

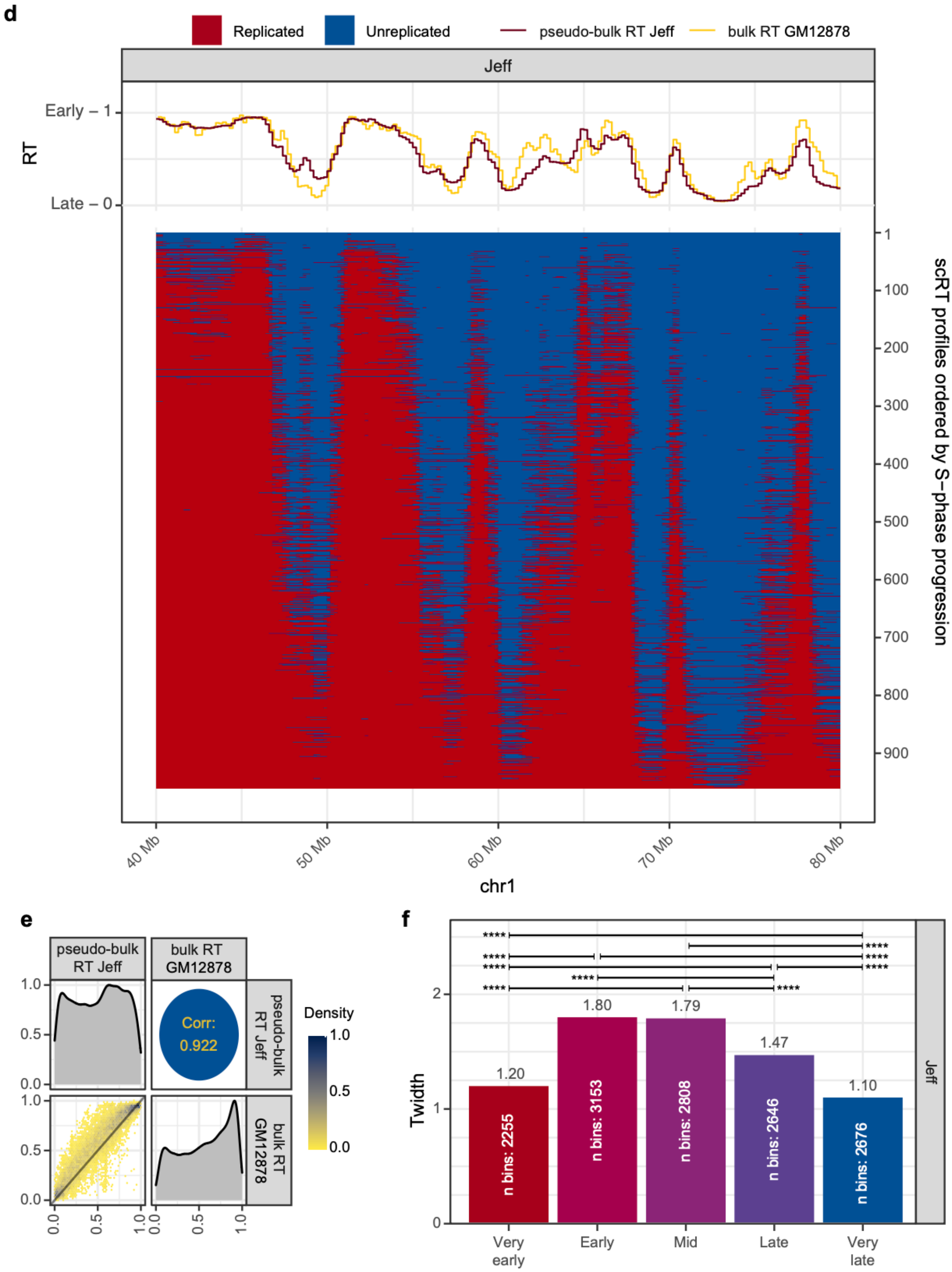


**Supplementary Figure 4. scRT from S-phase enriched human cells (extension of Fig. 5 for the HeLa and Jeff cells). a,d** The scRT of S-phase enriched HeLa (a) and Jeff (d) cells over a representative region. In the upper part of the plot, pseudo-bulk RT and bulk RT profiles of the correspondent cell line. In the bottom panel, the scRT profiles ordered from top to bottom by replication percentage of each cell. **b,e** Pairwise comparison of the HeLa (b) and Jeff (e) pseudo-bulk RT and bulk RT. Same as in

Supplementary Fig. 2. **c,f** Barplots reporting the $T_{widths}$ calculated on 5 RT categories based on the pseudo-bulk RT values in the HeLa (c) and Jeff (f). Categories were selected as in Fig. 3d. P-values were calculated using the Kronos scRT Compare TW module (Methods, * < 0.05, ** < 10-2, *** < 10-3, **** < 10-4).

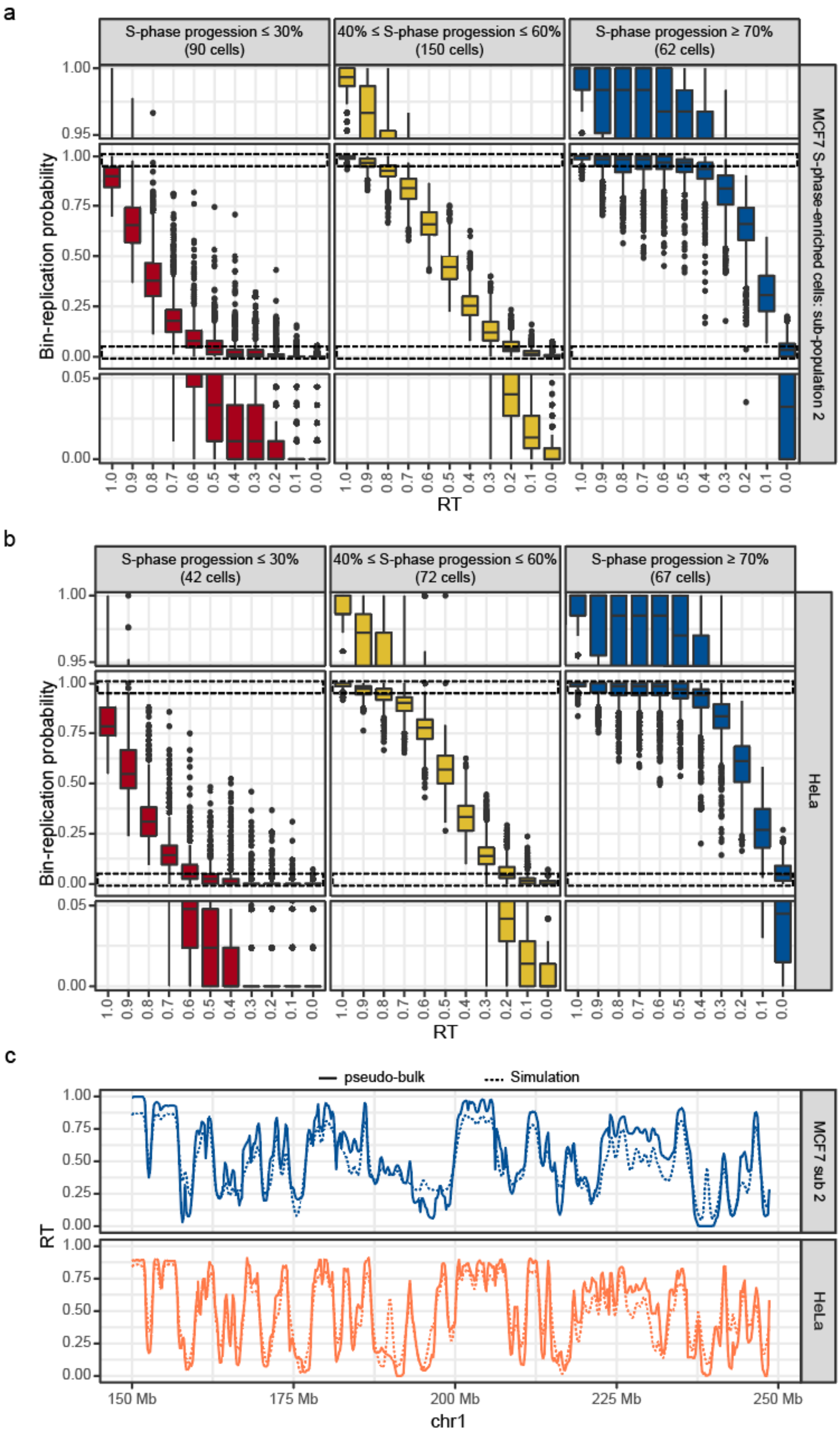
a
S-phase progression ≤ 30% (90 cells)
40% ≤ S-phase progression ≤ 60% (150 cells)
S-phase progression ≥ 70% (62 cells)
MCF7 S-phase-enriched cells: sub-population 2
Bin-replication probability
RT
b
S-phase progression ≤ 30% (42 cells)
40% ≤ S-phase progression ≤ 60% (72 cells)
S-phase progression ≥ 70% (67 cells)
HeLa
Bin-replication probability
RT
c
pseudo-bulk
Simulation
MCF7 sub 2
HeLa
RT
150 Mb
175 Mb
200 Mb
225 Mb
250 Mb
chr1

d

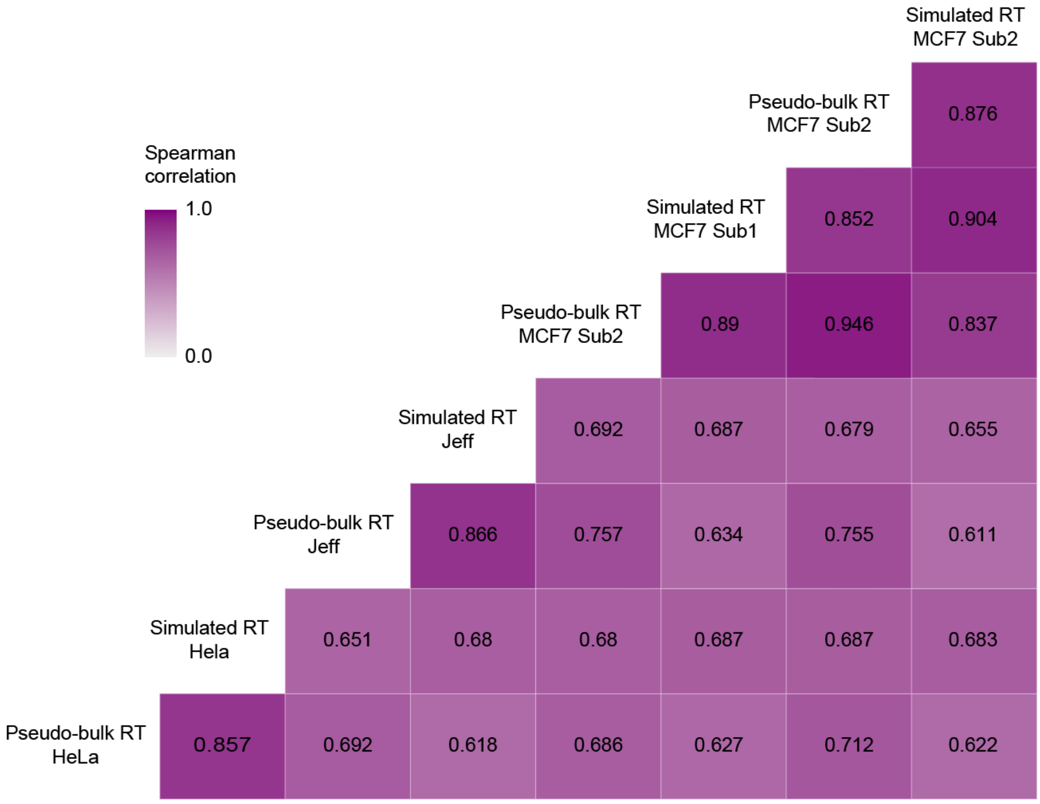


**Supplementary Figure 5. scRT data support a stochastic model of replication (extension of Fig. 6). a-b** Boxplots reporting the probability of a bin being replicated (y-axis) relative to its pseudo-bulk RT (x-axis, 1 is early and 0 is late) for S-phase-enriched MCF7 sub-population 2 (a) and HeLa (b) cells at different S-phase stages. The middle panel of each plot shows the whole distribution, and the top and bottom panels are zoom in for the extremities of the distributions (indicated with the dashed boxes in the middle panel). **c** Comparison between the pseudo-bulk RT (solid line) and simulated RT (dashed line) for the MCF7 sub-population 2 and HeLa cells. The simulation is based on Replicon[29] and uses the probability of being replicated within early-S-phase cells (completed up to 30% of their genome replication) for each 200 kb bin as input. **d** Pairwise Spearman correlation of the pseudo-bulk RT and simulated RT profiles for all the samples used in this study.
